# Discrete LAT condensates encode antigen information from single pMHC:TCR binding events

**DOI:** 10.1101/2021.12.16.472676

**Authors:** Darren B. McAffee, Mark K. O’Dair, Jenny J. Lin, Shalini T. Low-Nam, Kiera B. Wilhelm, Sungi Kim, Shumpei Morita, Jay T. Groves

**Affiliations:** Department of Chemistry, University of California, Berkeley, CA 94720, USA; Department of Chemistry, Purdue University, West Lafayette, ID 47907, USA

## Abstract

LAT assembly into a two-dimensional protein condensate is a prominent feature of antigen discrimination by T cells. Here, we use single-molecule imaging techniques to resolve the spatial position and temporal duration of each pMHC:TCR molecular binding event while simultaneously monitoring LAT condensation at the membrane. An individual binding event is sufficient to trigger a LAT condensate, which is self-limiting, and neither its size nor lifetime is correlated with the duration of the originating pMHC:TCR binding event. Only the probability of the LAT condensate forming is related to the pMHC:TCR binding dwell time. LAT condenses abruptly, but after an extended delay from the originating binding event. A LAT mutation that facilitates phosphorylation at the PLC-γ1 recruitment site shortens the delay time to LAT condensation and alters T cell antigen specificity. These results identify a role for the LAT protein condensation phase transition in setting antigen discrimination thresholds in T cells.

## INTRODUCTION

A healthy adaptive immune response depends on the ability of T cells to discriminate between agonist peptide major histocompatibility complex (pMHC) ligands and the vastly more abundant self pMHC. It has long been recognized that antigen discrimination is based on the binding kinetics of pMHC to T cell receptor (TCR), especially the kinetic off-rate (or, equivalently, the mean binding dwell time) (Corr et al., 1994; Daniels et al., 2006; Matsui et al., 1994). This task is complicated by the fact that agonist and self-ligands may differ only slightly in their binding kinetics, requiring a highly precise discrimination mechanism (Au-Yeung et al., 2014; François and Altan-Bonnet, 2016; Kersh et al., 1998; Lyons et al., 1996; McKeithan, 1995; Naeher et al., 2007). Furthermore, T cells can accurately discriminate pMHC ligands based on only a small number (tens) of individual molecular binding events (Huang et al., 2013; Irvine et al., 2002; Lin et al., 2019; O’Donoghue et al., 2013; Pielak et al., 2017). A quantitative mapping of the antigen discrimination function for an individual TCR has yet to be directly measured, and estimates based on indirect bulk parameters span a wide range (Chakraborty and Weiss, 2014; Jönsson et al., 2016; Merwe and Dushek, 2011; Pettmann et al., 2021; Stepanek et al., 2014). Nevertheless, precise tuning of the T cell signaling system is crucial for a proper immune response. Mutations in early signaling proteins that increase TCR sensitivity are associated with autoimmune diseases, while mutations that decrease TCR sensitivity are associated with immunodeficiency (Au-Yeung et al., 2018).

Binding of pMHC to TCR initiates a signaling process, the first few steps of which rely on sustained engagement of the pMHC:TCR complex (Alam et al., 1996; Chakraborty and Weiss, 2014; Matsui et al., 1994). This establishes a kinetic proofreading mechanism, in which only sufficiently long dwelling pMHC:TCR binding events successfully trigger the full downstream signaling response (Ganti et al., 2020; McKeithan, 1995; Merwe and Dushek, 2011). Following the initial formation of the pMHC:TCR complex, immune receptor tyrosine-based activation motifs (ITAMs) on the TCR CD3 chains are phosphorylated by the Src family kinase, Lck, creating docking sites for the Syk kinase, ZAP-70 (Chakraborty and Weiss, 2014; Iwashima et al., 1994). ZAP-70 arrives to the TCR in an initially autoinhibited state, but is activated after phosphorylation by Lck, and subsequently phosphorylates substrates including linker for activation of T cell (LAT) (Deindl et al., 2007; Klammt et al., 2015; Wang et al., 2010; Zhang et al., 1998). A distinctive feature of the TCR signaling mechanism is that LAT is a poor substrate for most kinases (Shah et al., 2016), and thus relies primarily on ZAP-70. This requisite series of kinase activation steps leading up to ZAP-70 activation at the TCR forms the first stage of kinetic antigen discrimination.

Under the control of phosphorylation by ZAP-70, LAT scaffolds a signaling hub downstream of TCR from which both Ca^2+^ and MAPK signal pathways branch (Balagopalan et al., 2015; Brownlie and Zamoyska, 2013; Finco et al., 1998; Samelson, 2002). LAT is an intrinsically disordered protein anchored to the membrane via a single transmembrane domain. LAT contains 9 tyrosine phosphorylation sites, at least 4 of which (Y136, Y175, Y195, Y235 in murine LAT) are utilized in T cell signaling. The Src-homology 2 (SH2) domain-containing adaptor protein Grb2 binds to phosphorylated LAT residues and subsequently recruits the Ras guanine nucleotide exchange factor (GEF), Son of Sevenless (SOS), which activates Ras and the MAPK pathway. Phosphorylation at tyrosine 136 on murine LAT (Y132 on human LAT) generates a binding site selective for PLC-γ1 recruitment (Houtman et al., 2004), which further interacts with GADS and SLP76, and ultimately activates Ca^2+^ signaling (Balagopalan et al., 2015). MAPK and Ca^2+^ signaling in T cells leads to ERK and NFAT nuclear translocation, respectively, and ultimately controls IL-2 production, T cell differentiation, and other effector functions (Smith-Garvin et al., 2009). It has long been known that the high multivalency of LAT phosphotyrosine sites, along with several crosslinking interactions among the various adaptor proteins, enables extended clustering into an elaborate signaling complex (Houtman et al., 2006; Kortum et al., 2013; Nag et al., 2009). More recent studies of LAT, Grb2, and SOS reconstituted on supported membranes have revealed a two-dimensional protein condensation phase transition governed by tyrosine phosphorylation (Huang et al., 2016; Su et al., 2016). Formation of this condensate was further discovered to facilitate release of autoinhibition in SOS, thus enabling Ras activation, and possibly providing a signal gating function in T cells (Huang et al., 2019).

In the work described here, we examine the signaling process from initial binding of individual pMHC:TCR molecular complexes to the associated LAT condensation. T cells are highly sensitive to antigen and can robustly activate at agonist pMHC densities around 0.2 molecules μm^-2^ (Irvine et al., 2002; Lin et al., 2019; Manz et al., 2011). At these densities, pMHC ligands are spaced microns apart and can be readily resolved by single-molecule imaging. We utilize the hybrid live cell—supported membrane system, in which a supported membrane functionalized with pMHC and intercellular adhesion molecule-1 (ICAM-1) forms a surrogate antigen presenting cell (APC) surface for interactions with primary mouse T cells. This experimental platform has long been used in studies of T cell signaling in the context of the immunological synapse (Campi et al., 2005; Chang et al., 2016; Grakoui et al., 1999; Huppa et al., 2010; Ma et al., 2017; Schmid et al., 2016; Taylor et al., 2017; Zhao et al., 2018), and provides an optimal configuration for single-molecule imaging by total internal reflection fluorescence (TIRF) microscopy (Douglass and Vale, 2005; Huppa et al., 2010; O’Donoghue et al., 2013). Individual pMHC molecules in the supported membrane can be tracked with high spatial (50 nm) and temporal (20 ms) resolution. In these experiments, free pMHC molecules exhibit simple two-dimensional Brownian motion in the membrane and nonspecifically immobilized pMHC represent a negligibly small fraction (< 1%). When pMHC binds TCR on an opposed T cell, however, its motion changes dramatically, allowing clear distinction of pMHC:TCR complexes from free pMHC ligand (Lin et al., 2019; O’Donoghue et al., 2013). Here, we use this single-molecule imaging strategy to track the formation, duration, and movement of individual pMHC:TCR complexes while simultaneously monitoring LAT condensation (as well as localization of other signaling molecules) in response to each pMHC:TCR binding event. Unique to this experimental strategy is the ability to map the LAT condensation and other signaling responses to the specific molecular binding event from which they originated.

The observations reveal that a single pMHC:TCR binding event is sufficient to trigger formation of a two-dimensional condensate on the membrane containing hundreds of LAT molecules. LAT condensation occurs abruptly, but after an extended delay from the originating binding event, exhibiting signatures of a phase transition. The resulting LAT condensates are self-limiting, and neither their size nor their lifetime is correlated with the duration of the originating pMHC:TCR binding event. Only the probability of forming a LAT condensate is related to the pMHC:TCR binding dwell time. We report quantitative measurements of this probability distribution, providing experimental mapping of a single TCR antigen discrimination function. These results reveal extended kinetic discrimination of ligand, into the tens of seconds binding dwell times. This long timescale discrimination represents an additional layer of kinetic proof-reading, extending beyond the TCR itself, which is provided by the LAT condensation phase transition. We further observe that a LAT mutation (G135D in mouse, G131D in human), which enhances the kinetics of ZAP-70 phosphorylation at Y136 on LAT (the PLC-γ1 recruitment site) (Lo et al., 2019), decreases the delay time to LAT condensation and alters T cell antigen specificity. Whole cell activation (measured by NFAT translocation) correlates with the number of LAT condensates formed after exposure to pMHC, suggesting that LAT condensates represent quanta of information in the T cell signaling pathway.

Collectively, these results reveal a central role for the LAT condensation phase transition in translating antigen information, from individual pMHC:TCR binding events, to downstream signaling pathways. The ensuing LAT condensates contain hundreds of LAT molecules and establish significant amplification from the single originating pMHC ligand. Additionally, the self-limiting characteristics of the resultant condensate discretize information into a binary output. Individually, a pMHC:TCR binding event either produces a LAT condensate or it doesn’t, but once formed, all condensates function independently from the originating binding event. We suggest that the LAT condensation phase transition has evolved in TCR signaling as means to achieve the amplification and noise suppression necessary for the single-molecule ligand sensitivity exhibited by T cells.

## RESULTS

### Single pMHC:TCR binding events trigger discrete LAT condensates

We characterize LAT condensation in response to pMHC:TCR binding events using a hybrid live cell-supported membrane experimental platform (Biswas and Groves, 2019; Brian and McConnell, 1984; Céspedes et al., 2021; Grakoui et al., 1999; Mossman et al., 2005) (**Figure 1A**). The supported membrane, consisting primarily of DOPC lipids, is functionalized with pMHC (0 − 100 molecules μm^−2^) and ICAM-1 (300 − 600 molecules μm^−2^), both of which are linked to the membrane via His-tag-protein:Ni-chelating-lipid interactions (Groves and Dustin, 2003; Nye and Groves, 2008). In the experiments described here, we utilize multicolor TIRF imaging to track LAT condensation at the membrane in response to T cell engagement with the pMHC and ICAM-1 functionalized supported bilayers. Experiments are generally performed using primary T cells harvested from mice with transgenic TCR(AND) (Kaye et al., 1989). Splenocytes from the TCR(AND) mice, hemizygous for H2^k^, were pulsed with 1 μM moth cytochrome c (MCC) peptide and cultured with the T cells for two days. The T cell blasts were treated with IL-2 from the day after harvest to the fifth day after harvest, at which point the cells were used in experiments. Fluorescent fusion proteins of interest were expressed using the PLAT-E retroviral platform (Morita et al., 2000); for experiments requiring expression of two different fluorescent fusion proteins we utilized a self-cleaving P2A peptide (Kim et al., 2011). Reflection interference contrast microscopy (RICM) was used as a way of monitoring the cell-supported membrane contact independently from the fluorescence channels (Limozin and Sengupta, 2009).

**Figure 1.**
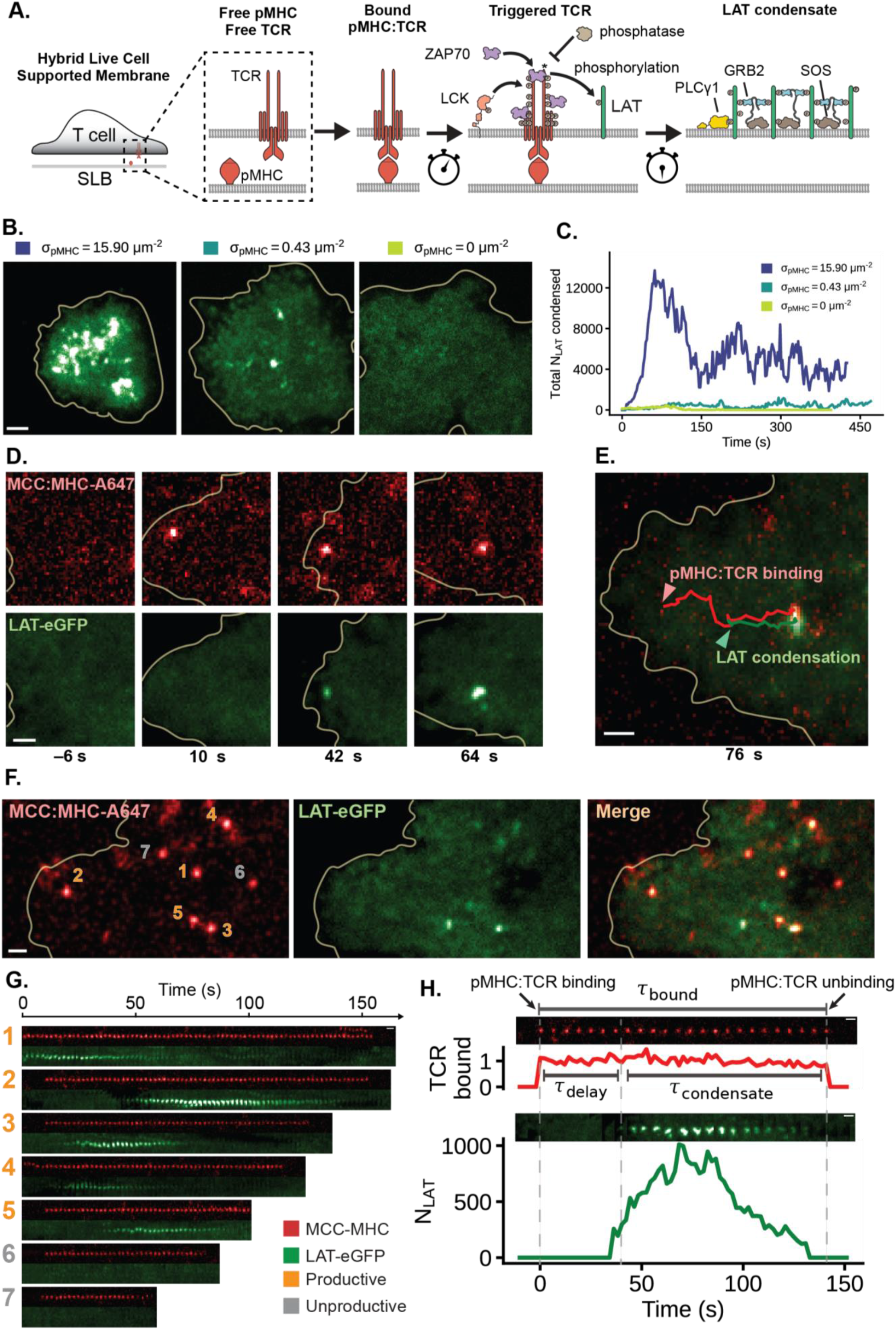
Single pMHC:TCR binding events trigger discrete LAT condensates. (A) Schematic of T cell signaling that occurs downstream of pMHC:TCR binding after the plasma membrane of the T cell interfaces with the supported lipid bilayer (SLB). (B) TIRF images of primary murine CD4+ T cells expressing AND TCR and LAT-eGFP deposited onto SLBs with varying densities of MCC(Atto647) pMHC. (C) The amount of condensed LAT through time for cells in B. and was calculated by dividing the intensity of the total condensed area (as determined by simple thresholding) by the intensity of single LAT-eGFP molecules, see Methods for more detail. (D) Time series images of a spatially isolated pMHC:TCR binding event that produces a highly localized LAT condensate within 50 nm at onset of condensation. Time stamps are relative to the moment of pMHC:TCR binding. (E) Overlay of MCC(Atto647) pMHC and LAT-eGFP channels showing the co-localized trajectories of the pMHC:TCR binding event and its associated LAT condensate, as well as their centripetal motion towards the center of the cell. (F) A wider view of the T cell shows a constellation of binding events spread out in space. Separate binding events are enumerated. (G) Temporal sequence of images for the binding events in F. Binding events visible in the first frame of the data acquisition are not considered for subsequent analyses. Some binding events are productive, while others fail to produce a localized LAT condensate. (H) Intensity traces of productive binding events have several quantities - the number of condensed LAT produced (*N*_*LAT*_), the delay time until LAT condensation (*τ*_delay_), as well as the lifetime of the LAT condensate (*τ*_condensate_). Related data are in Figure S1.

TIRF microscopy of the live cell-supported membrane interface provides robust imaging capabilities all the way down to the single molecule level (Douglass and Vale, 2005; Jaiswal and Simon, 2007; Zlatanova and Holde, 2006). At low pMHC densities (0.1 − 0.4 molecules μm^−2^) individual pMHC molecules can be tracked undergoing free Brownian motion laterally in the membrane, with diffusion coefficients of 0.55 ± 0.07 μm^2^s^−1^ as determined from single-molecule step size distributions. When pMHC binds to a TCR on a T cell, its motion changes dramatically to conform with the much slower movement of the TCR (**Movie S1, Figure S1A**) (DeMond et al., 2008; Lin et al., 2019; O’Donoghue et al., 2013; Yu et al., 2012). Under long exposure times (500 ms) and low excitation powers (0.4 mW), individual pMHC:TCR complexes appear as well defined spots whereas free pMHC is moving too quickly to be clearly imaged, and appears as a diffuse background (**Figure S1B**). This imaging method offers selective tracking of pMHC:TCR complexes in living cells with time resolution of about one second and spatial resolution of ≈ 300 nm (Lin et al., 2019; O’Donoghue et al., 2013). Trajectories for each pMHC:TCR complex are tracked to measure the specific pMHC:TCR binding dwell time as well as their spatial movement within the T cell membrane.

Representative TIRF images of primary T cells expressing LAT-eGFP interacting with stimulatory bilayers containing various densities of agonist pMHC are illustrated in **Figure 1B** (see also **Movie S2**). LAT condensation is readily visible as localized increases in LAT density and corresponding time traces of the overall extent of LAT condensation are plotted in **Figure 1C**. At high agonist pMHC densities of 10 − 40 molecules μm^−2^, LAT condensation is observed throughout the cell-supported membrane interface within seconds of cell landing. This is consistent with numerous reports of LAT clustering and condensation in T cells interacting with APCs displaying high agonist pMHC density (Balagopalan et al., 2018; Campi et al., 2005; Huse et al., 2007; Muhammad et al., 2009) as well as T cells interacting with activating anti-TCR antibody coated surfaces (Bunnell et al., 2002; Yi et al., 2019). At these high antigen densities LAT phosphorylation and subsequent condensation is driven by multiple TCR activation events, which are also intrinsically un-synchronized in time and, are superimposed in the images. The observed LAT condensation is a conglomerate of correspondingly unsynchronized individual nucleation events (**Figure S1C**), which cannot be directly mapped to the specific pMHC:TCR binding events that triggered them; and much information is lost. At lower agonist pMHC densities (**Figure 1B**, middle panel, also see **Movie S3**), the overall amount of LAT condensation is lower, however similarly high local densities of LAT are still observed to form stochastically and widely distributed throughout the interface. Although the total amount of LAT condensation remains low, and appears to develop on a much longer timescale, T cells still activate robustly at these pMHC densities (Huang et al., 2013; Irvine et al., 2002; Lin et al., 2019; Manz et al., 2011). In control experiments with zero pMHC in the membrane (**Figure 1B**, right panel), very few LAT condensates are observed and T cells do not activate. The different timescales observed for LAT condensation at low and high antigen are in good agreement when the superposition of intrinsically stochastic individual binding events is taken into consideration (see *Simulation of Whole Cell Integrated LAT Signal* in Methods and **Figures S1C, S1D, S1E**).

At low pMHC densities (0.1 − 0.4 molecules μm^−2^), individual pMHC:TCR binding events are spaced microns apart and can remain distinct for the entire lifetime of the bound complex. Individual pMHC:TCR binding events can nucleate LAT condensation. A time sequence of images illustrating the formation and movement of a single pMHC:TCR complex along with the corresponding LAT condensate it nucleated are illustrated in **Figure 1D** (see also **Movie S4**). LAT condensation begins within tens of nanometers of the originating pMHC:TCR complex and generally remains within a 100 − 300 nm neighborhood of the complex as both undergo colocalized transport towards the center of the cell (**Figure 1E** and **S1F**). This retrograde transport of single pMHC:TCR complexes along with their associated LAT condensate is very similar to the retrograde transport of TCR clusters that occurs at high pMHC density (DeMond et al., 2008; Dustin and Groves, 2012; Yokosuka et al., 2005; Yu et al., 2012). LAT condensates are observed to nucleate throughout the cell-supported membrane interface, with a slight preference for more peripheral positions (**Figure S1G**). The imaging methods we use here are also sensitive enough to distinguish rarely formed pMHC dimers (**Figure S1H**) from the predominant monomer pMHC:TCR complexes; we focus on monomer pMHC:TCR binding events at low overall agonist pMHC densities in this study. Over the entire cell interface, multiple such pMHC:TCR binding events and associated LAT condensates can be observed (**Figure 1F; Movie S5**). The LAT condensates form and ultimately dissipate, generally independently of each other; at any one moment in time there may only exist a few LAT condensates within the cell. A collection of image time sequences tracking pMHC:TCR binding along with the associated LAT condensate formation and subsequent dissipation for the seven binding events from the cell pictured in **Figure 1F** are shown in **Figure 1G**; note that two binding events failed to produce LAT condensates in this particular set.

Detailed analysis of a productive binding event is presented in **Figure 1H**, illustrating the information that is gathered from each such event. The total pMHC:TCR binding dwell time (*τ*_bound_) is observed in the top (red) sequence of images. The sequence of images below (green) track the corresponding LAT condensate, with a calibrated trace plotting the number of LAT molecules in the condensate through time. LAT condensation occurs abruptly, but after a relatively long delay time (≈ 40 sec. in this trace; mean delay is ⟨*τ*_delay_⟩ ≈ 23 seconds). The LAT condensates are self-limiting and, as seen here, can dissipate prior to the pMHC:TCR complex dissociating. Fluorescence recovery after photobleaching measurements on the LAT condensates (**Figure S2A**) indicate they are relatively dynamic, with individual LAT molecules turning over throughout the lifetime of the condensate (the effective off-rate for LAT within the condensate was measured at *k*_off_ = 0.24 *s*^−1^; though this is likely a lower-bound estimate; see *FRAP of LAT condensates* in Methods).

We determined the total number of LAT molecules in each pMHC:TCR-induced condensate using a combination of quantitative immunoblots (**Figure S2B**), fluorescence activated cell sorting (FACS), and quantitative fluorescence imaging (**Figure S2C, Movie S7**, see *Quantitative Fluorescence* in Methods). The measured fluorescence intensity within a LAT condensate relative to the local background level in the surrounding membrane (**Figure 2A**) reveals the relative increase in LAT density in the condensate. Overall expression levels of endogenous LAT and exogenous LAT-eGFP were determined through quantitative immunoblots and FACS. The observed LAT copy number in single condensates was measured to be a linear function of total LAT expression level (**Figure 2B**). By extrapolating to endogenous LAT expression levels (zero expression of LAT-eGFP), we determine that 258 ± 65 (SD) LAT molecules are contained within a physiological condensate produced by a single pMHC:TCR binding event. In **Figure 2C** we plot data from an ensemble (n = 100) of pMHC:TCR binding events mapping the binding dwell time, which contains the physical information of antigen identity, to properties of the associated LAT condensate. Neither the size, nor the lifetime, of a LAT condensate is correlated with the duration of the originating pMHC:TCR binding event. As analyzed in greater detail below, only the probability of LAT condensation formation is related to pMHC:TCR binding dwell time. Other characteristics of the LAT condensate do not contain antigen information.

**Figure 2.**
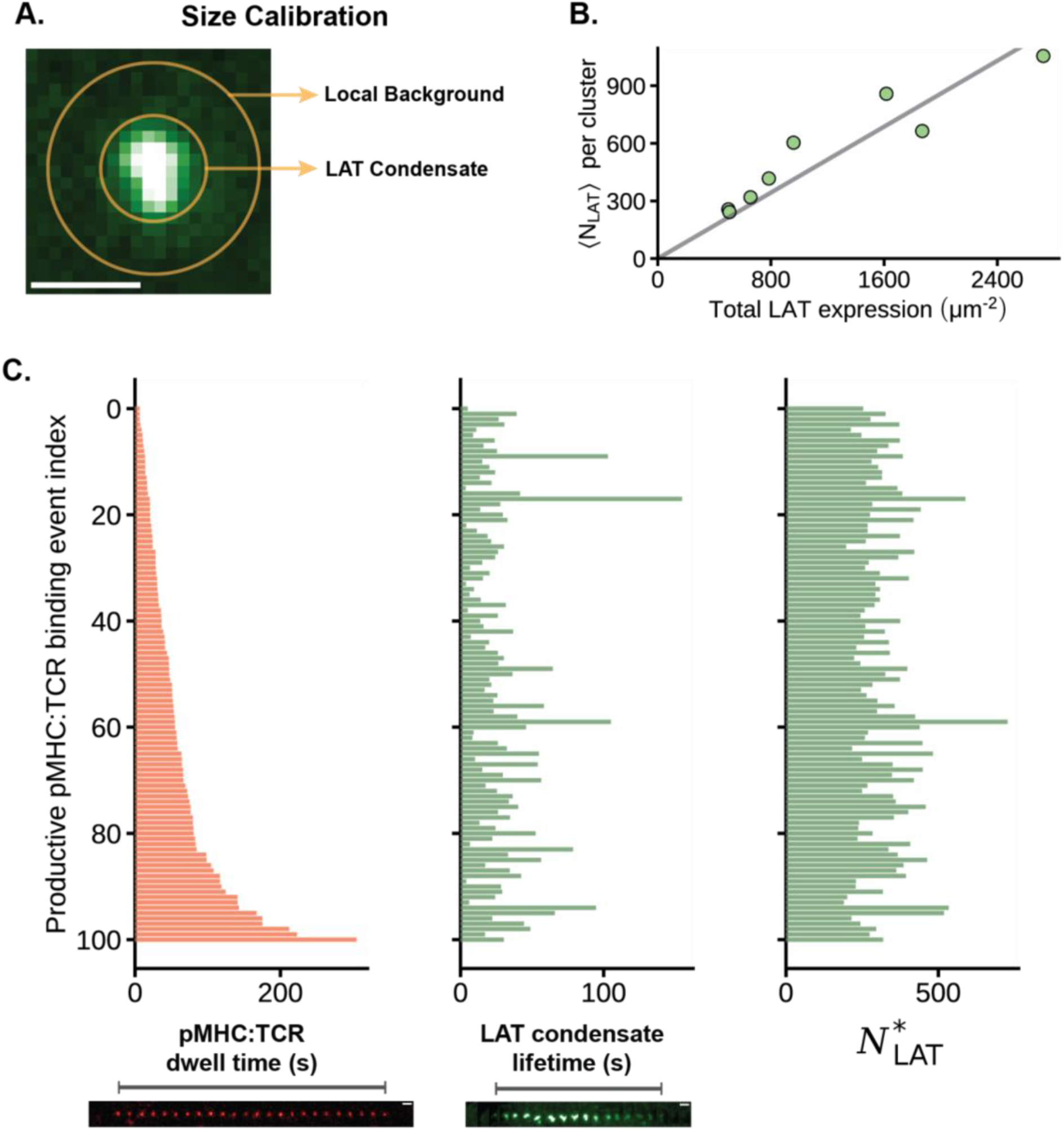
Physical properties of the LAT condensate. (A) An annulus around the LAT condensate is used to sample the local background. The intensity of the condensate is calculated as the midpoint of the net intensity and total intensity, since the extent of background LAT clustering is unknown. (B) Scatter plot of the average number of LAT within condensates, ⟨*N*_*LAT*_⟩, as a function of the total LAT expression level (endogenous LAT + exogenous LAT-eGFP). Each point is a cell average over condensates observed within the first 5 minutes of cell landing. The fitted line has the equation *y* = 0.43 *x*, which was constrained to have 0 intercept. All cells were from the same mouse. Cells from two other mice (data not shown), yielded similar fits, *y* = 0.51*x* and *y* = 0.48*x*. (C) Three bar plots showing the lack of correlation between pMHC:TCR dwell time (left) with either the LAT condensate lifetime (middle) or the number of LAT within its associated condensate (right). Each red bar is a pMHC:TCR binding event, data is pooled from the 8 cells in B. The relative size of the LAT condensate is calculated as 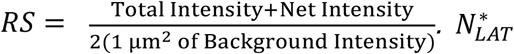 is the relative size (RS) of the condensate rescaled using an estimated physiological (zero exogenous) LAT density of 601 ± 150 (SD) *μm*^−2^. This normalizes LAT cluster size to LAT expression level for cell-to-cell comparisons. Related data are in Figure S2.

The background distribution of LAT in primary T cell membranes is extremely uniform (as seen in **Figures 1B** and **1D**). Individual pMHC:TCR binding events produce distinctly resolved LAT condensates against this smooth background. Experimental examination reveals that these pMHC:TCR-induced LAT condensates form from plasma membrane LAT and are distinct, in both composition and their association with agonist pMHC, from LAT vesicles arriving at the membrane from the cytosol (a small number of which can also be observed). Delivery of cytosolic LAT to the plasma membrane has been reported via VAMP7-associated vesicles (Balagopalan et al., 2018; Soares et al., 2013). To confirm the pMHC:TCR-induced condensates studied here are distinct, we performed experiments on T cells simultaneously expressing LAT-mScarleti and VAMP7-mNeonGreen. LAT condensates associated with pMHC:TCR binding events did not show any appreciable inclusion of VAMP7 (**Figure S2D**). However, at later time points (>30 s after cell landing), some dense features of LAT did appear in the TIRF images that were not associated with pMHC:TCR and contained VAMP7 (**Figure S2E**). Some LAT features were also observed that lacked VAMP7 and were not associated with detectable pMHC:TCR (**Figure S2F**), possibly corresponding to pMHC:TCR-induced condensates in which the fluorescent label on pMHC has been bleached or some other form of LAT oligomerization (Raab et al., 2017). In this study we focus on LAT condensates whose origin can be directly associated to a pMHC:TCR binding event. We also note that Jurkat T cells can exhibit greater variability and can have significant levels of LAT pre-clustering or pre-condensation in the absence of any TCR activation (**Figure S2G**, top), rendering them less suitable for these types of high-resolution single-molecule studies. Control experiments looking at primary human T cells revealed negligible pre-clustered LAT, while stimulation with anti-CD3 antibody confirmed clean LAT condensation events exist (**Figure S2G**, bottom). Thus, the above-mentioned issues are specific to the immortalized human Jurkat T cell line, and do not represent a difference between mouse and human primary T cells.

### pMHC:TCR binding dwell time modulates probability of LAT condensate formation

A single pMHC:TCR binding event either produces a local LAT condensate or fails to do so (**Figure S3A**). Although the size and lifetime of individual LAT condensates exhibit no correlation with the binding dwell time of the originating pMHC:TCR complex (**Figure 2C**), the probability of producing a condensate is correlated. Here we define the success probability, *P*_*LAT*_(*τ*_*MHC*_), to be the probability that a pMHC:TCR binding event of duration *τ*_*MHC*_ produces a LAT condensate at any point during its lifetime. We measure this success probability function for an individual TCR by examining an ensemble (n=1071) of individual MCC pMHC:TCR binding events and mapping the corresponding LAT condensation outcome. Data from this ensemble is compiled in **Figure 3A**, illustrating the pMHC:TCR binding dwell time (traced in red) as well as subsequent LAT condensation (traced in green), if it occurs. The delay time between initial pMHC:TCR binding and LAT condensation (*τ*_*delay*_) is also visible in this data set, but here we focus exclusively on the binary success or failure of each event. The data is aggregated into bins based on pMHC:TCR dwell time, indicated by grey lines, and the resulting counts of all and successful binding events are plotted in **Figure 3B**. A similar data set derived from the weaker T102S pMHC is plotted in **Figure 3C**. The observed dwell time distributions from all binding events for both the strong (MCC) and moderate (T102S) agonist pMHC:TCR are exponentially distributed with mean dwell times of 23.8 ± 2.4 (SE) and 9.1 ± 2.8 (SE) seconds, respectively. These observed dwell times include events that end by unbinding as well as photobleaching, but imaging parameters are adjusted to minimize photobleaching (Lin et al., 2019) while maximizing temporal resolution of the moment of LAT condensation (see *Imaging* in Methods). These *in situ* dwell time measurements, when combined with photobleaching measurements, allow extraction of the true binding dwell times of 44 ± 6 (MCC) and 10.8 ± 2 (T102S) seconds (see *Dwell Times* in Methods), which are in good agreement with bulk kinetic measurements for pMHC:TCR binding for MCC and T102S (Corse et al., 2010; Newell et al., 2011). In both cases, productive binding events exhibited longer mean dwell times than the population average (52.2 ± 6.1 (SE) and 15.6 ± 3.3 (SE) seconds for MCC and T102S, respectively). The success probability as a function of dwell time, *P*_*LAT*_(*τ*_*MHC*_), based on the time bins plotted in **Figures 3B** and **3C**, is plotted in **Figure 3D** (bottom). Since a binding event is considered successful if a LAT condensate forms at any point during its lifetime, this distribution is a calibrated measure of probability; it is not impacted by the sampling efficiency for different binding dwell times (see *P*_*LAT*_(*τ*_*MHC*_) *and Delay Time* in Methods). These data reveal that short dwelling binding events have near zero probability of producing a LAT condensate, but as binding dwell time increases, so does the probability of LAT condensation, with a *t*_half−max_ = 24.0 ± 1.3 seconds. The longest dwelling events approach a constant 0.25 probability of producing a LAT condensate. The similarity of the probability curves traced by both MCC and T102S pMHC reinforces the primacy of dwell time over peptide identity as the fundamental input to TCR triggering, at least between these two peptide antigens.

**Figure 3.**
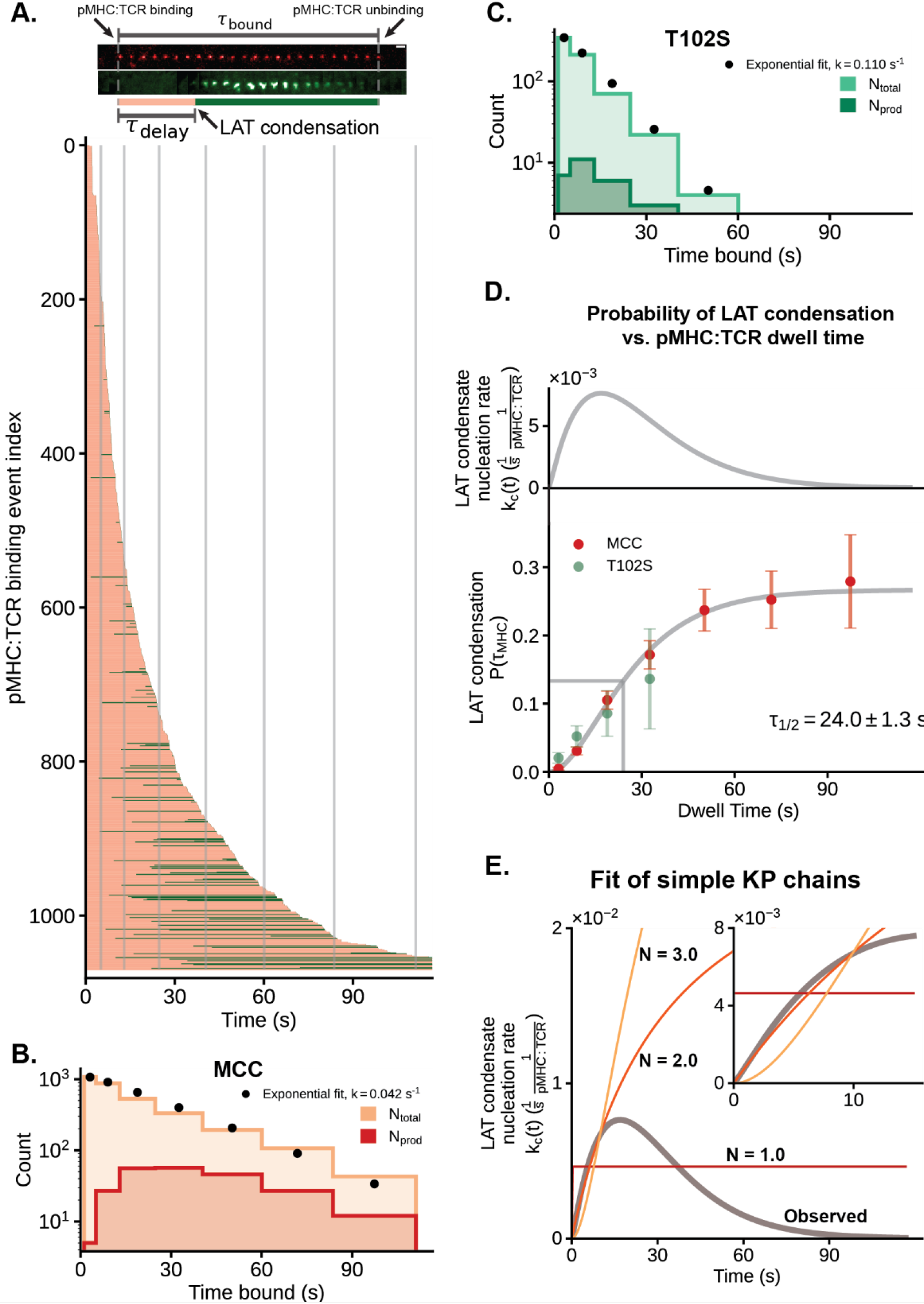
pMHC:TCR binding dwell time modulates probability of LAT condensate formation. (A) Top: time series images illustrating the temporal partitioning of the total pMHC:TCR dwell time into a delay time (pink) and a remaining dwell time (green). Bottom: A bar plot of a collection of 1071 pMHC:TCR binding events from 12 cells across 3 mice. Each bar represents a fully tracked pMHC:TCR binding event. The moment of condensation (if any) is indicated by the bar transitioning from pink (unproductive dwell time) to green (productive dwell time). The gray lines demarcate the bins used in the histograms for B. and C. (B) MCC pMHC dwell time segments were aggregated according to the indicated bin widths. The number of productive dwell time segments (red) for a particular time bin was the number binding events within that dwell time window which had at some prior time produced a LAT condensate. The total population of dwell time segments (orange) was fit with an exponential decay rate of 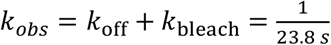. The fit distribution was integrated for each bin and plotted as a black circle. (C) T102S pMHC dwell time segments were aggregated according to the indicated bin widths. The number of productive dwell time segments (green) for a particular time bin was the number binding events within that dwell time window which had at some prior time produced a LAT condensate. The total population of dwell time segments (aquamarine) was fit with an exponential decay rate of 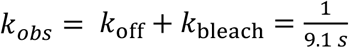. The fit distribution was integrated for each bin and plotted as a black circle. (D) Bottom: The probability of a pMHC:TCR binding event producing a localized LAT condensate as a function of dwell time. For each bin of the histograms in B and C, the fraction of dwell time segments that were productive is plotted as a point. The error bars are the standard error of the mean for a binomial variable. The interpolating gray line was a fitted regularized gamma function with a maximal amplitude parameter, *f*(*t*). Top: The effective rate of LAT production: 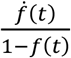 is plotted, where *f*(*t*) is the gray interpolating line from the bottom panel. (E) The effective nucleation rate for various gamma distributions were fit to the early rise in the observed nucleation rate. Related data in Figure S3.

The data plotted in **Figure 3D** represents measurement of a single TCR antigen discrimination function. It reveals two mechanistic features of molecular antigen discrimination by TCR not readily discerned from bulk measurements. First, the long rise time after initial binding indicates that individual TCR are still sensitive to the binding dwell time of a pMHC ligand beyond 30 seconds from initial engagement. This is much longer than kinetic cut-offs for antigen discrimination observed at high antigen density (1 – 3 s) (Bridgeman et al., 2012; Pettmann et al., 2021; Stepanek et al., 2014), and represents an additional layer of kinetic proofreading that becomes prominent when antigen counts are low. Although the measured probability of a pMHC:TCR complex inducing a LAT condensate after just a few seconds is small (0.05 within 10 seconds) it is not zero. Engagement of a sufficiently large number of TCR with pMHC, as occurs under high antigen exposure, will overcome this low probability step by sheer numbers. At low antigen densities, however, long dwelling pMHC:TCR binding events to 30 s and beyond become important—as has been observed experimentally in single-molecule T cell activation studies (Lin et al., 2019).

The second mechanistic discovery is that the probability function for an individual pMHC:TCR complex to induce a LAT condensate plateaus at a maximum of only 0.25. The fact that this success probability plateaus significantly below a value of one indicates the presence of some form of localized negative feedback specifically timed relative to each individual binding event. This is clearly revealed by examining the momentary effective rate constant, *k*_*c*_(*t*), for formation of a LAT condensate, which is the probability per unit time of a LAT condensation event occurring a time *t* after initial binding of pMHC to TCR. The measured success probability function, *P*_*LAT*_(*τ*_*MHC*_), is related to *k*_*c*_(*t*) through the following integral equation: 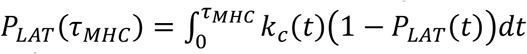, which on differentiation yields 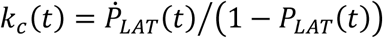. The functional form of *k*_*c*_(*t*) extracted from success probability measurements (**Figure 3D**, upper panel) exhibits an initial rising phase, over the first ≈ 15 s, then begins declining and reaches essentially zero after 60 – 90 s. Practically, this reveals that a pMHC:TCR complex can only produce a LAT condensate within the first minute or so of existence. Beyond this time, even if the TCR remains engaged with pMHC, there is essentially no chance of a condensate forming. This is a single receptor level observation. The T cell itself is not shut off, and newly formed pMHC:TCR can still produce LAT condensates following the same success probability function, with the clock started for each TCR at the moment it binds pMHC. Some plausible mechanisms for how this relative timing of localized negative feedback may be achieved in T cells include further LAT modification – e.g., ubiquitylation (Balagopalan et al., 2007) – or ZAP-70 inactivation (Sjölin-Goodfellow et al., 2015). Both of these processes would occur locally with timescales set by individual LAT condensation or initial pMHC:TCR binding time. Additionally, once formed, pMHC:TCR complexes begin to move in a directed retrograde flow pattern towards the center of the immunological synapse. Early studies mapping geometric position within the immunological synapse to TCR signaling activity identified differential effects in the immunological synapse center favoring signal downregulation (Lee et al., 2003; Mossman et al., 2005). It is plausible that the prevailing cytoplasmic conditions in more central positions also disfavor LAT condensation and moving LAT condensates into this region might facilitate dispersal.

### Comparison with classic kinetic proofreading schemes

Measurement of the single TCR antigen discrimination function, *P*_*LAT*_(*τ*_*MHC*_), and the corresponding *k*_*c*_(*t*), also provides a useful point for quantitative comparison with classic kinetic proofreading schemes. In its most basic form, kinetic proofreading for antigen discrimination by TCR has been considered as a sequence of transitions beginning with pMHC binding to TCR. The TCR doesn’t achieve activation until the final step and if pMHC unbinds before this point, activation is not achieved (Chakraborty and Weiss, 2014; McKeithan, 1995). If the intermediate steps have similar rates, the strength of kinetic proofreading—referring to how sharply defined is the transition between activating and nonactivating ligands—can be roughly measured by the number of steps. Several studies of TCR and related systems have sought to infer this effective number of steps from downstream signaling parameters measured over the whole cell, such as diacylglycerol production (Tischer and Weiner, 2019), Ca^2+^ flux (Yousefi et al., 2019) and IL-2/CD69 expression (Pettmann et al., 2021).

Kinetic proofreading processes are often examined in terms of a steady state rate of activation (François et al., 2013; McKeithan, 1995; Pettmann et al., 2021). However, this is a simplifying limit case and a more complete description of a kinetic proofreading mechanism is provided by the delay time distribution between initial ligand engagement of the receptor and subsequent activation, here referred to as *f*_*D*_(*t*). Note that if we define the associated LAT condensation as the full activation event for a TCR, then *f*_*D*_(*t*) is the derivative of the measured single TCR antigen discrimination function, 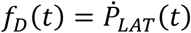. For a series of *N* intermediate steps, *f*_*D*_(*t*) = *f*_1_(*t*)⨂*f*_2_(*t*)⨂ … ⨂*f*_*N*_(*t*), where *f*_*i*_(*t*) are the individual delay time distributions at each step, and *f*_*D*_(*t*) results from their successive convolution. In the case where each step is a first order process with equivalent rate, *λ*, this reduces to a gamma distribution: 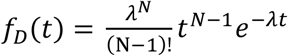, and can be used to calculate the momentary effective rate constant *k*_*c*_(*t*) (see *k*_*c*_(*t*) *as a Hazard Rate* in Methods). When there are no intermediate steps (*N* = 1), *k*_*c*_(*t*) is constant and there is no kinetic proofreading. For *N* greater than one, *k*_*c*_(*t*) is a rising function that asymptotically approaches a maximum value when kinetic proofreading is satisfied. At early time points (< 13 s), before substantial negative feedback has kicked in, the experimental data agrees well with a simple kinetic proofreading mechanism with *N* = 2.0 and an intermediate step rate of *λ* = 0.03 *s*^−1^, though this is not strongly distinguished from mechanisms with more steps (**Figure 3E**). The marked decline in observed *k*_*c*_(*t*) at longer times observed in the experimental data is not predicted by classic kinetic proofreading mechanisms.

### LAT condensation occurs after an extended delay

For productive binding events, LAT condensation occurs after a distinctively long delay from the originating pMHC:TCR binding event. When the LAT condensation begins, the growth rate rapidly transitions from zero to rates near 100 LAT molecules per second (see **Figures 1H** and **S4A**); and this growth is sustained as hundreds of LAT molecules join the condensate, reaching a maximum after 9.3 ± 6 (SD) seconds (**Figure S4B**). The abrupt transition to LAT condensation establishes a well-defined delay time (*τ*_delay_) between the originating pMHC:TCR binding event and LAT condensation (**Figure 4A**). A histogram of 115 delay times measured from productive MCC pMHC:TCR binding events reveals a broad distribution, with the rise and fall shape of a gamma distribution, and a mean delay time of 23.2 ± 1.6 (SE) seconds (**Figure 4B**). These measurements have a time resolution of 2 seconds, which is set by the time-lapse used in the image sequences. A collection of 91 trajectories of productive pMHC:TCR binding events can be visualized in **Figure S4C**. Control experiments confirm that the identity of the fluorophore on the LAT has no effect on LAT condensation (see LAT-mCherry condensation in **Movie S6**) nor was the delay time affected by LAT expression levels (**Figure S4F**).

**Figure 4.**
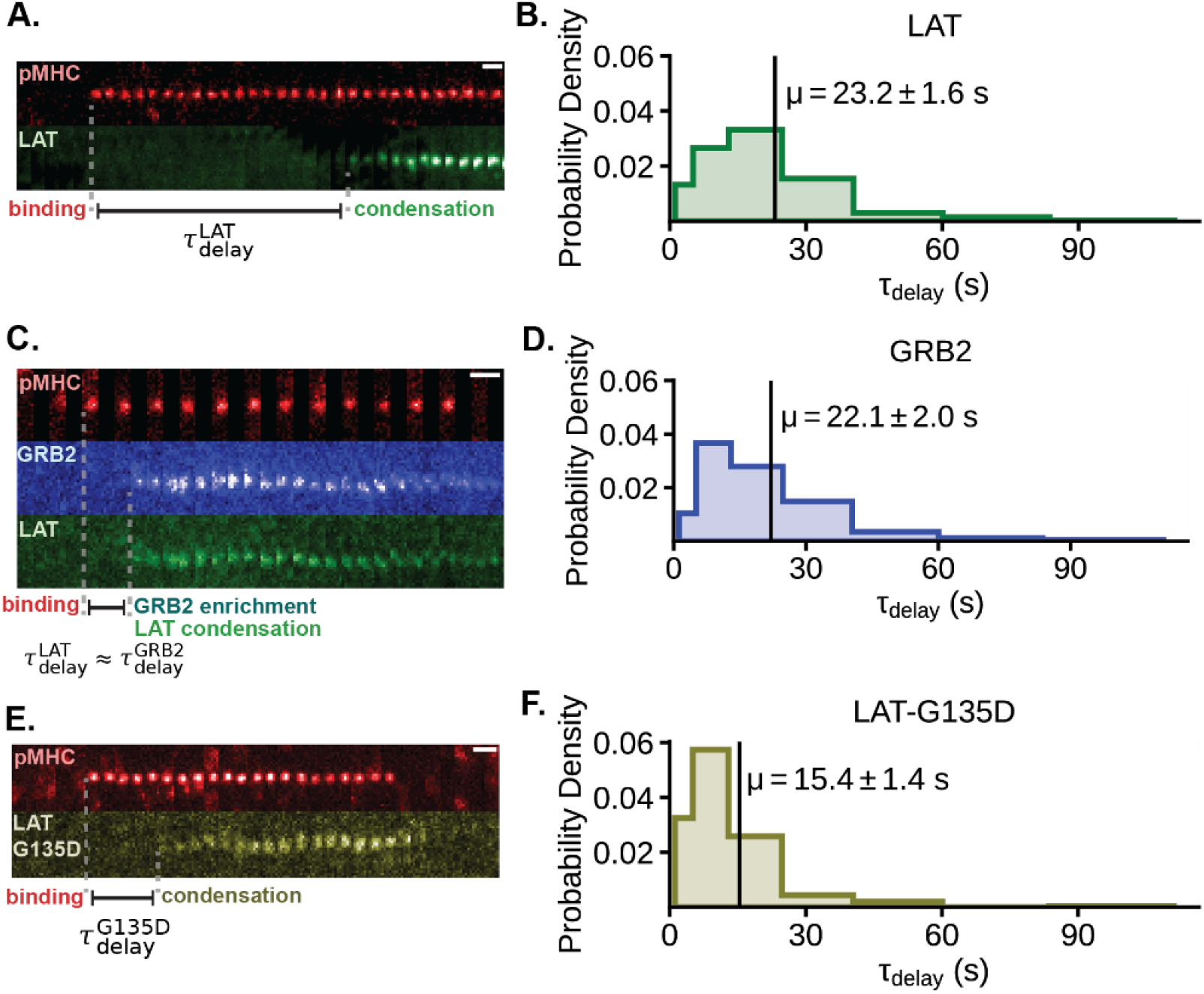
LAT condensation occurs after an extended delay. T cells expressing different constructs were deposited onto bilayers with 0.1 − 0.2 molecules μm^−2^ of MCC(Atto647) pMHC. (A) TIRF images of LAT-eGFP condensation within primary murine T cells in response to a single pMHC:TCR binding event. LAT condensation occurs near the binding event after a long delay, 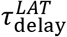, measured relative to the moment of pMHC:TCR binding. (B) Histogram of an ensemble LAT condensate delay times (115 binding events, from 18 cells across 4 mice). (C) TIRF images of LAT-mScarleti and mNeonGreen-GRB2 clustering within primary murine T cells in response to a single pMHC:TCR binding event. Delay times of LAT clustering, 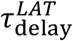, and GRB2 clustering, 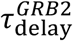, relative to binding were equivalent within the resolution of this experiment (< 2 s). (D) Histogram of an ensemble GRB2 clustering delay times from primary murine T cells expressing only mNeonGreen-GRB2 (75 binding events, from 21 cells across 4 mice). (E) TIRF images of LAT(G135D)-eGFP condensation within primary murine T cells in response to a single pMHC:TCR binding event. (F) Histogram of an ensemble of delay times between pMHC:TCR binding and LAT(G135D) condensation (102 binding events, from 17 cells across 3 mice). Related data are in Figure S4.

We next address the question of what is causing the delay to LAT condensation by examining the localized concentration of phosphorylated LAT prior to condensation. Grb2 binds several phosphorylated tyrosine residues on LAT (primarily Y175, Y195, and Y235 in mouse), and does so independently of LAT condensation. Grb2 has been used as a precision probe for detailed kinetic studies of LAT phosphorylation (Huang et al., 2016, 2017) and here we used Grb2 to monitor LAT phosphorylation in T cells. In this experiment, primary AND T cells were transduced with a bicistronic P2A vector containing LAT-mScarleti and mNeonGreen-GRB2. TIRF imaging experiments were performed as described before, but now simultaneously monitoring single pMHC:TCR binding events, LAT condensation and Grb2 recruitment in three channels. Enrichment of Grb2 was always observed simultaneously with LAT condensation with generally no detectable sustained Grb2 enrichment prior to condensation (**Figure 4C** and **S4D**). To rule out potential artifacts from incomplete cleavage of the P2A peptide between the two proteins, we performed further experiments with a LAT(4F)-mCherry and mNeonGreen-GRB2 linked by P2A peptide. The LAT(4F) mutant has all three primary Grb2 tyrosine binding sites (Y175, Y195, Y235) as well as the PLC-γ1 site (Y136) mutated to phenylalanine. The mutant LAT(4F) failed to participate in any condensates while the simultaneously expressed Grb2 was observed to condense with endogenous LAT(WT) (**Figure S4E**). As an additional control experiment, monocistronic mNeonGreen-GRB2 was also used to compile a histogram of Grb2 enrichment delay times, which provide an alternative measure of LAT condensation (**Figure 4D**). The resulting distribution, with a mean delay time of 22.1 ± 2.0 (SE) seconds, is nearly identical to the measured distribution using LAT imaging as the readout (**Figure 4B**).

These results indicate that densities of sustained phosphorylated LAT remain undetectable above background prior to condensation. Competition from phosphatases dephosphorylating LAT and diffusion of LAT away from the active pMHC:TCR complex are likely responsible for limiting localized accumulation of phosphorylated LAT. This observation rules out one possible cause of the delay between pMHC:TCR binding and LAT condensation: that it takes a sustained period of ZAP-70 kinase activity to build up a sufficient density of phosphorylated LAT before the phase transition can occur. Rather, it appears that a composition fluctuation from the competing kinase-phosphatase reactions themselves is the nucleating event. This conclusion is further supported by the broad distribution of measured delay times; some LAT condensates form within a few seconds of the originating pMHC:TCR binding event while others begin to form after one minute.

### Phosphorylation kinetics at the PLC-γ1 binding site on LAT control condensation delay time

Although the delay time to LAT condensation is insensitive to both LAT and Grb2 expression levels, we find that a LAT point mutation that modulates phosphorylation kinetics by ZAP-70 substantially reduces the delay time. In human wild type LAT, the glycine immediately upstream of Y132, the PLC-γ1 binding site, makes Y132 a poor substrate for ZAP-70 compared to Y171 and Y191. By substituting the glycine for a negatively charged aspartate, the rate of Y132 phosphorylation by ZAP-70 dramatically increases (Shah et al., 2016). We performed the homologous mutation in murine LAT, G135D, in order to enhance the phosphorylation rate of Y136. Primary T cells expressing LAT(G135D)-eGFP over a background of endogenous wild type LAT continued to exhibit LAT condensation events (observed by their incorporation of the labeled LAT(G135D) mutant) in response to single pMHC:TCR binding events (**Figure 4E**). However, the delay time between pMHC:TCR binding and LAT(G135D) condensation was dramatically reduced (**Figure 4F**). The resulting distribution, with a mean delay time of 15.4 ± 1.4 (SE) seconds, was narrower than that of LAT(WT); and the acceleration was achieved despite the presence of large amounts of endogenous wild-type LAT.

### The relation between LAT condensates and T cell activation

LAT condensation (referred to as clustering in older publications) has long been recognized as a critical element of T cell activation by TCR (Houtman et al., 2006; Kortum et al., 2013; Lin and Weiss, 2001). However, most prior work has examined bulk LAT condensation in response to high antigen exposure, where hundreds to thousands of times more TCR are activated than are necessary to trigger a full T cell response (Crites et al., 2014; Huse et al., 2007; Su et al., 2016; Yi et al., 2019). Here we examine the low antigen end of the spectrum to measure how the sparsely distributed LAT condensates stemming from individual pMHC:TCR binding events relate to T cell activation. We first examine LAT condensation in response to a panel of altered peptides (MCC, T102S, ER60, and T102E), spanning a range of mean dwell times and corresponding potencies (Pielak et al., 2017; Schild et al., 1995). T cells were exposed to supported membranes containing 600 molecules μm^−2^ of ICAM-1 and 0.5 molecules μm^−2^ pMHC after loading MHC with one of the peptides from the panel. In most conditions, high amounts of the null T102E pMHC (10 molecules μm^−2^) were included as background for the tested pMHC to mimic the abundant self pMHC that exists on antigen presenting cells *in vivo*. LAT condensation events were tracked over time by TIRF imaging using the LAT-eGFP construct (**Figure 5A**). The null T102E pMHC + ICAM-1 condition itself did not produce any LAT signal over background levels observed with ICAM-1 only conditions. After 200 seconds, the longer dwelling MCC peptide (*τ*_dwell_ ≈ 45 seconds) produced 83 ± 35 (SD) distinct LAT condensates, while the shorter dwelling agonist T102S (*τ*_dwell_ ≈ 10 seconds) and non-agonist ER60 (*τ*_dwell_ ≈ 1 seconds) produced 42 ± 20 (SD) and 8 ± 2 (SD) LAT condensates, respectively. This trend is generally consistent with known potencies for these pMHC (Crites et al., 2014).

**Figure 5.**
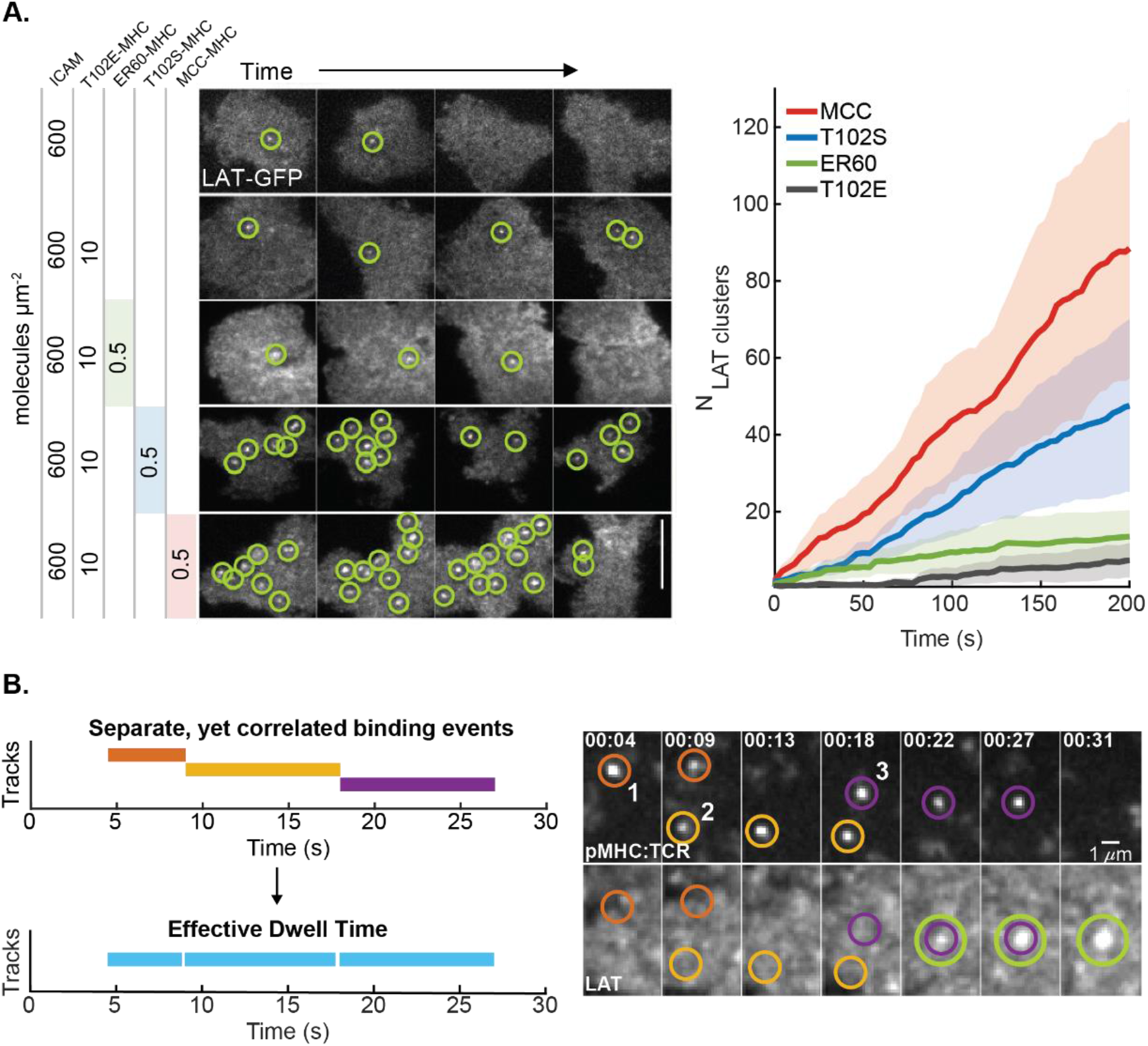
The number of LAT condensates correlates with ligand potency. (A) Left: a time series of TIRF images of primary murine T cells deposited onto supported membranes containing 600 molecules μm^−2^ ICAM-1 and one of four distinct peptides at the indicated concentrations: MCC(Atto647), T102S(Atto647), ER60(Atto647), or T102E. Right: cumulative time-series for the number of distinct LAT condensates observed. From most productive to least: MCC (red), T102S (blue), ER60 (green), and lastly T102E (gray). Compare to Figure S2G. (B) Left: schematic of how distinct binding events, that are locally coincident, may potentiate LAT condensation. Right: TIRF images showing rapid LAT condensation after a series a locally coincident T102S(Atto647) pMHC:TCR binding events. Related data are in Figure S5.

In the cases of both MCC and T102S, more LAT condensation events were observed over the entire cell than simple extrapolation from single TCR measurements would predict. Some of these extra LAT condensates represent inclusion of other types of LAT clusters not directly produced from pMHC:TCR binding events (e.g., VAMP-7 containing LAT vesicles as noted above) in these measurements. However, even when other LAT clusters and potential rebinding of pMHC are considered, T102S pMHC stimulation still produces disproportionately more LAT condensates than MCC pMHC (see *Extrapolation from P*(*τ*_*pMHC*_) in Methods and **Figure S5A** and **S5B**). Close examination of the T102S pMHC:TCR binding events and associated LAT condensates reveals a type of correlation that may be responsible for the apparently enhanced sensitivity of T cells to the shorter-dwelling T102S pMHC ligand. **Figure 5B** illustrates an example of spatiotemporally correlated T102S pMHC:TCR binding events that appear to nucleate a single LAT condensate. Evidence of such spatiotemporal correlations enhancing the potency of short dwell time ligands has been detected previously in experiments mapping pMHC:TCR binding events to T cell activation (Lin et al., 2019). Such effects may also have contributed to earlier observations of apparent pMHC:TCR cooperativity at pMHC levels so low that multiple pMHC:TCR complexes were unlikely to coexist at the same time (Manz et al., 2011). Here we present visualization data suggesting that the LAT condensation phase transition itself is an integrating mechanism capable of establishing cooperativity between pMHC:TCR complexes that do not exist simultaneously. We further note that ER60 pMHC, which has approximately a fifth of the binding dwell time with TCR as T102S pMHC, exhibited very little signal over null peptide conditions. This suggests there exists a hard cutoff around 1 second pMHC:TCR binding dwell time for downstream signaling, which is in good agreement with other measurements (Bridgeman et al., 2012; Stepanek et al., 2014).

To quantify the relation between LAT condensation and T cell activation, we simultaneously monitored the formation of LAT condensates and NFAT translocation in response to pMHC:TCR binding events (**Figure 6A**). NFAT translocation to the nucleus occurs downstream of Ca^2+^ activation in T cells and has been successfully used as a read-out of early T cell activation (Marangoni et al., 2013; Melichar et al., 2013; Podtschaske et al., 2007). Here, we expand on an assay previously developed by our lab that mapped patterns of pMHC:TCR binding to NFAT translocation (Lin et al., 2019) by incorporating LAT-eGFP into the expression vector using a P2A peptide. In this manner, we can simultaneously monitor T cell adhesion, pMHC:TCR binding, LAT condensation, and the cellular activation state in real time. For these experiments, T cells were incubated on bilayers with ICAM-1 and either MCC pMHC or T102S pMHC at 0.22 molecules μm^−2^ and were monitored for 12 minutes, after which we measured the final activation state (**Figure 6B**). Even though a smaller proportion of cells activated in response to T102S pMHC, those cells which did activate experienced a similar number of LAT condensates as cells activating in response to MCC pMHC (**Figure 6C**). This data indicates that it takes 45 − 75 LAT condensation events within 12 minutes to generate sufficient signal to induce Ca^2+^ flux and subsequent NFAT translocation. The actual minimum number of LAT condensates may be lower due to the lag time of NFAT translocation.

**Figure 6.**
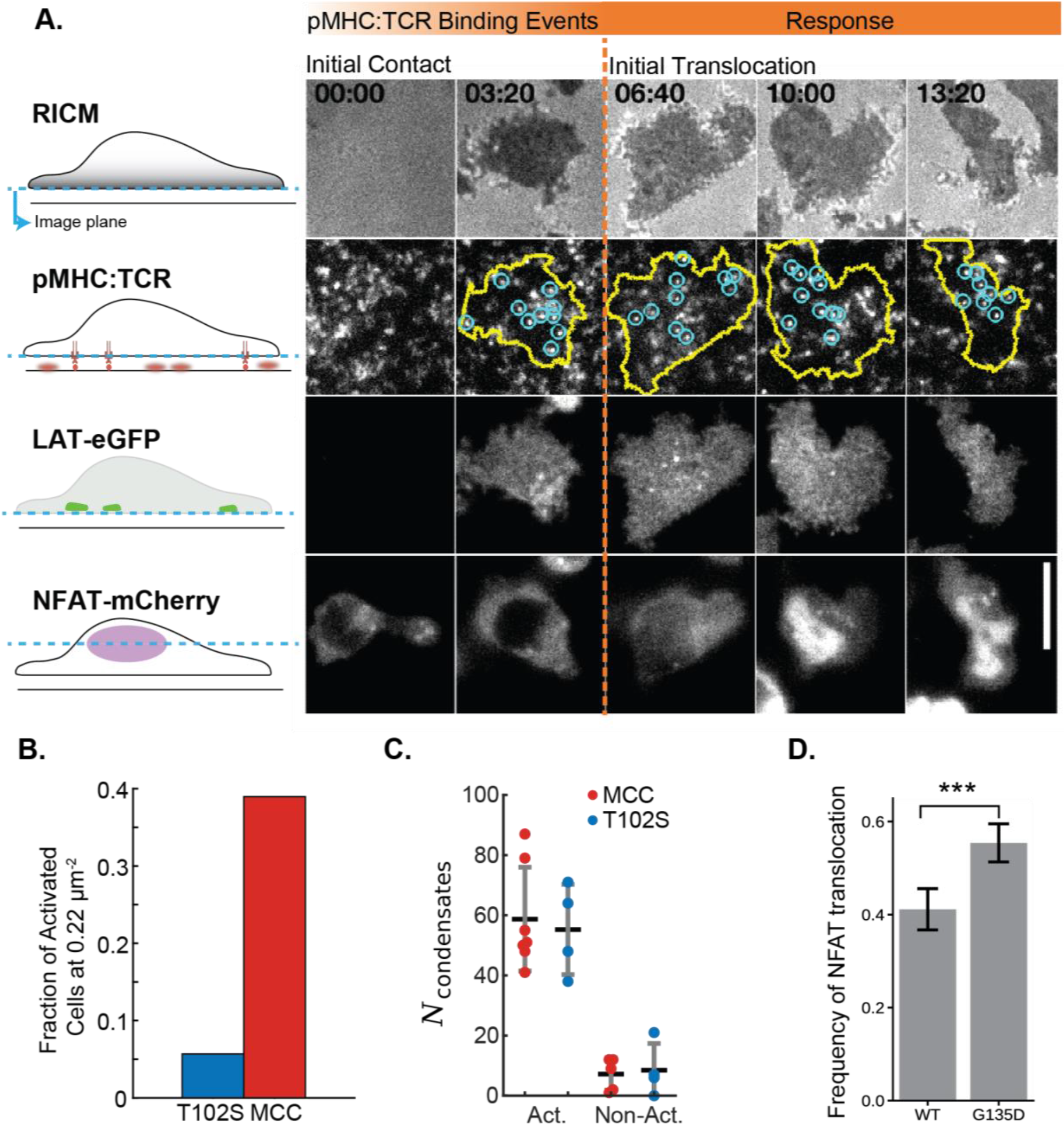
Modulating LAT condensate delay time alters antigen discrimination thresholds. (A) Images of key molecular steps leading to NFAT translocation through time within primary T cells. The cells were deposited onto supported membranes containing ICAM-1 and MCC(Atto647) pMHC. First row, RICM detects cell spreading and ensures the T cell has good contact with the supported membrane. Second row, TIRF images of individual pMHC:TCR binding events (blue) are recorded within the cell perimeter (yellow). Third row, TIRF images of LAT-eGFP is monitored for condensation events. Fourth row, epifluorescence images of NFAT to track moment of translocation. (B) Bar plot showing the frequency of T cells that undergo NFAT translocation for either T102S or MCC at a fixed peptide-MHC density of 0.22 molecules μm^−2^. (C) Plot showing the number of distinct LAT condensates formed prior to NFAT translocation (Act.) or the number of LAT condensates formed prior to 300 s if no translocation was observed (Non-Act.) (D) Bar plot showing the frequency of NFAT translocation for T cells on T102S-MHC bilayers (at 1.1 molecules μm^−2^) after 30 minutes as a function of the LAT construct expressed, either WT or G135D.

Finally, we assess how the delay time between LAT condensation and pMHC:TCR binding affects antigen discrimination using the LAT(G135D) mutant. The G135D mutation has been shown to enhance both Ca^2+^ and ERK activity in response to weaker agonists, particularly at low stimulation thresholds (Lo et al., 2019). Results comparing NFAT translocation rates between T cells expressing LAT(WT) or LAT(G135D) reveal that the presence of LAT(G135D) increased NFAT translocation rates by ≈ 15% (**Figure 6D; Movie S8**). These experiments were done using the weaker T102S pMHC agonist at its activation threshold density, 1.1 molecules μm^−2^ (Lin et al., 2019). While the observed effect is modest, it provides important evidence that alterations in LAT condensation kinetics, which occur on the 20-30 second timescale, still change the T cell responsiveness to significantly shorter dwelling pMHC ligands.

## DISCUSSION

Certain output parameters of the T cell activation response, including NFAT and Erk activation as well as cytokine production, exhibit essentially binary behavior at the single cell level (Das et al., 2009; Huang et al., 2013; Podtschaske et al., 2007). When observing these parameters, exposure to higher antigen pMHC densities may increase the probability of an individual cell activating—or, equivalently, an apparently stronger activation response from a population of T cells—but it does not change signal strength at the single cell level. By contrast, many molecular intermediates in the TCR signaling pathway, including TCR phosphorylation, ZAP-70 recruitment, LAT phosphorylation, and LAT condensation exhibit a much more linear dose-response behavior with respect to the copy number of agonist pMHC molecules at moderate to high levels (Grakoui et al., 1999; Lo et al., 2018; Yokosuka et al., 2005). Titrating down to single molecule levels, TCR and ZAP-70 activation remain essentially stoichiometric with agonist pMHC (O’Donoghue et al., 2013) whereas LAT exhibits substantially different behavior. The stochastic formation of discrete LAT condensates (containing hundreds of LAT molecules) in response to individual pMHC:TCR binding events that we report here represents a distinct transition in the propagation of signal downstream from TCR. This reveals not only a major amplification step, but also a thresholding mechanism that quenches downstream signaling from shorter-lived, but still potentially ZAP-70 kinase active, pMHC:TCR complexes. The further observation that a LAT condensate’s size and lifetime are not correlated with the binding dwell time of its originating pMHC:TCR complex is indicative of a discretization mechanism, by which a continuum of pMHC:TCR binding dwell times is parsed into a binary success or failure output. This signal digitization would apply to signals dependent on formation of the LAT signaling complex; it is also possible some TCR signaling activity is independent of this – such as tonic signaling activity from very short-lived self pMHC:TCR interactions (Oers et al., 1993, 1994; Persaud et al., 2014).

Clustering and assembly of molecules in the TCR signaling pathway has been a topic of intense interest, from both an immunological (Grakoui et al., 1999; Yi et al., 2019; Yokosuka et al., 2005) and biophysical perspective (Hu et al., 2013; Merwe and Dushek, 2011; Mossman et al., 2005; Qi et al., 2001; Weikl and Lipowsky, 2004). Foundational experiments on LAT have mapped the presence of multiple distinct Grb2, GADS, and PLC-γ1 phosphotyrosine binding sites as required elements for downstream propagation of signaling from activated TCR (Hartgroves et al., 2003; Houtman et al., 2006; Zhang et al., 2000). Key realizations from this early work are that multiple pY sites needed to be on the same LAT molecule (e.g., multivalency is required) and that all these molecules condense into a LAT-scaffolded signaling microcluster (here referred to as a LAT condensate) in T cells (Balagopalan et al., 2015; Lin and Weiss, 2001). These early studies were almost invariably done with high agonist pMHC stimulation or antibody stimulation (which involves artificial TCR crosslinking). Here we have presented experimental results that zoom in on individual pMHC:TCR binding events for a live, superresolution view of the initiation of LAT condensation in T cells.

The single-molecule imaging experiments reveal signature behavior of a protein condensation phase transition that is not detectable in bulk observations. We observe that discrete LAT condensates form in response to individual pMHC:TCR binding events and that LAT condensation occurs abruptly, and after a well-defined delay time from the originating pMHC:TCR binding event. The delay times are broadly distributed, with some LAT condensates forming within a few seconds of the pMHC:TCR binding event while others may delay up to a minute. In all cases, there is no detectable build-up of localized pLAT prior to condensation. Collectively, these observations are consistent with a phase transition nucleation process controlled by fluctuations in phosphorylation reaction kinetics; the competing kinase-phosphatase reactions controlling LAT phosphorylation provide the nucleating event (Shimobayashi et al., 2021). These features of the phase transition are readily visible at low agonist pMHC densities because individual binding events remain well separated and can be unambiguously tracked for extended times. At higher pMHC densities, multiple LAT condensates originating from different pMHC:TCR binding events appear superimposed and unsynchronized in the composite LAT condensation response (**Figures 1B, 1C** and SI). This creates an illusion of a more continuous growth process while the underlying mechanism is likely to still involve the discrete phase transition we observe at the single molecule level.

A critical remaining question is what are the qualitative differences between a kinetic nucleation phase transition process and a more linear growth process with respect to signal propagation through LAT? Two key observations provide a clue: *i*) a single activated pMHC:TCR complex is not able to build up a localized concentration of pLAT prior to condensation; and *ii*) LAT condenses abruptly once nucleation is achieved. This combination of features reveals an initially strong resistance to signal propagation (e.g., LAT phosphorylation) followed by positive feedback that favors LAT phosphorylation once condensation has been initiated. This type of initial resistance followed by positive feedback is an effective noise filtering mechanism (commonly found in electronic devices such as avalanche photodiodes used for single photon detection (Saleh and Teich, 2019)). If LAT condensation were to follow a more linear phosphorylation and growth process at the single molecule level, this type of noise filtering mechanism would be defeated. A second insight is provided by recent studies on Ras activation by SOS in LAT condensates that identified a strong enhancement of SOS activity in the condensed state (Huang et al., 2019). Mechanistically, multivalency in the condensate retains SOS at the membrane for extended periods of time, providing sufficient time for autoinhibition release. It is plausible that a similar mechanism may govern PLC-γ1 activation.

Finally, the discrete and delayed condensation of LAT tied to individual pMHC:TCR binding events provides an additional layer of kinetic proofreading for antigen discrimination. This is revealed in our measurement of the single TCR antigen discrimination function (**Figure 3D**). The long mean delay time between LAT condensation and the originating pMHC:TCR binding event renders short dwelling pMHC:TCR complexes highly unlikely to produce a LAT condensate and any downstream signals that rely on this. This effect is most pronounced at low pMHC levels, where cooperativity between multiple pMHC:TCR complexes is essentially impossible since they are so widely spaced. At higher antigen densities, we have observed instances of a spatially localized sequence of short pMHC:TCR binding events evidently contributing to a single LAT condensate (**Figure 5B**). This reveals a cooperative mechanism that enables the T cell to respond to shorter dwelling pMHC ligands that are present at sufficiently high density. Effectively, the final stage of kinetic proofreading provided by the LAT condensation event is bypassed at higher antigen density since multiple pMHC:TCR may work together to achieve nucleation. Earlier stages of kinetic proofreading (e.g., Lck phosphorylation of ITAMS, ZAP-70 recruitment, and ZAP-70 activation) are still necessary and pMHC ligands that fail to pass these stages won’t activate T cells at any density (Corr et al., 1994; Matsui et al., 1994; Pettmann et al., 2021; Stepanek et al., 2014).

Protein condensation phase transitions are emerging as important mechanistic features in a wide range of biological processes (Hyman et al., 2014; Klosin et al., 2020; Patel et al., 2015; Shimobayashi et al., 2021; Su et al., 2016). Here, using single-molecule imaging methods, we have resolved signature characteristics of the LAT protein condensation phase transition in T cell activation. The results provide quantitative mapping of early signal filtering and amplification mechanisms in TCR signaling. More broadly, they reveal a number of functional capabilities that are provided by the phase transition process that may also be found in other membrane signaling systems.

## AUTHOR CONTRIBUTIONS

Conceptualization: J.T.G. and D.B.M.; Methodology: D.B.M.; Formal Analysis: D.B.M.; Investigation: D.B.M., M.K.O., S.T.L.-N., J.J.L., K.B.W., and S.M; Resources: D.B.M, J.J.L., and S.K.; Data Curation: D.B.M., M.K.O., S.T.L.-N., and J.J.L.; Writing – Original Draft: D.B.M; J.T.G; Writing – Review and Editing: D.B.M., M.K.O., S.T.L.-N., J.J.L., K.B.W., S.K, S.M., J.T.G.; Supervision: J.T.G.; Funding Acquisition: J.T.G.

## ACKNOWLEDGEMENTS

We thank S. Hansen (University of Oregon) and S. Alvarez (Inscripta) for their contributions to the synthesis of p2a plasmids and cloning advice used in this study; A. Weiss (UCSF) for gifting a sample of Jurkat cells; F. Marangoni (Harvard Medical School) for providing the NFAT reporter plasmid; L. Teyton (Scripps Research Institute) and M. Davis (Stanford University) for providing the MHC and ICAM-1 bacmids; and Kole T. Roybal for the primary human T cells. The financial support for this work was provided by National Institutes of Health grant P01 AI091580 and by the Novo Nordisk Foundation Challenge Programme as part of the Center for Geometrically Engineered Cellular Systems.

## DECLERATIONS OF INTEREST

We have no conflicting interests to declare.

## SUPPLEMENTAL FIGURES

**Figure S1.**
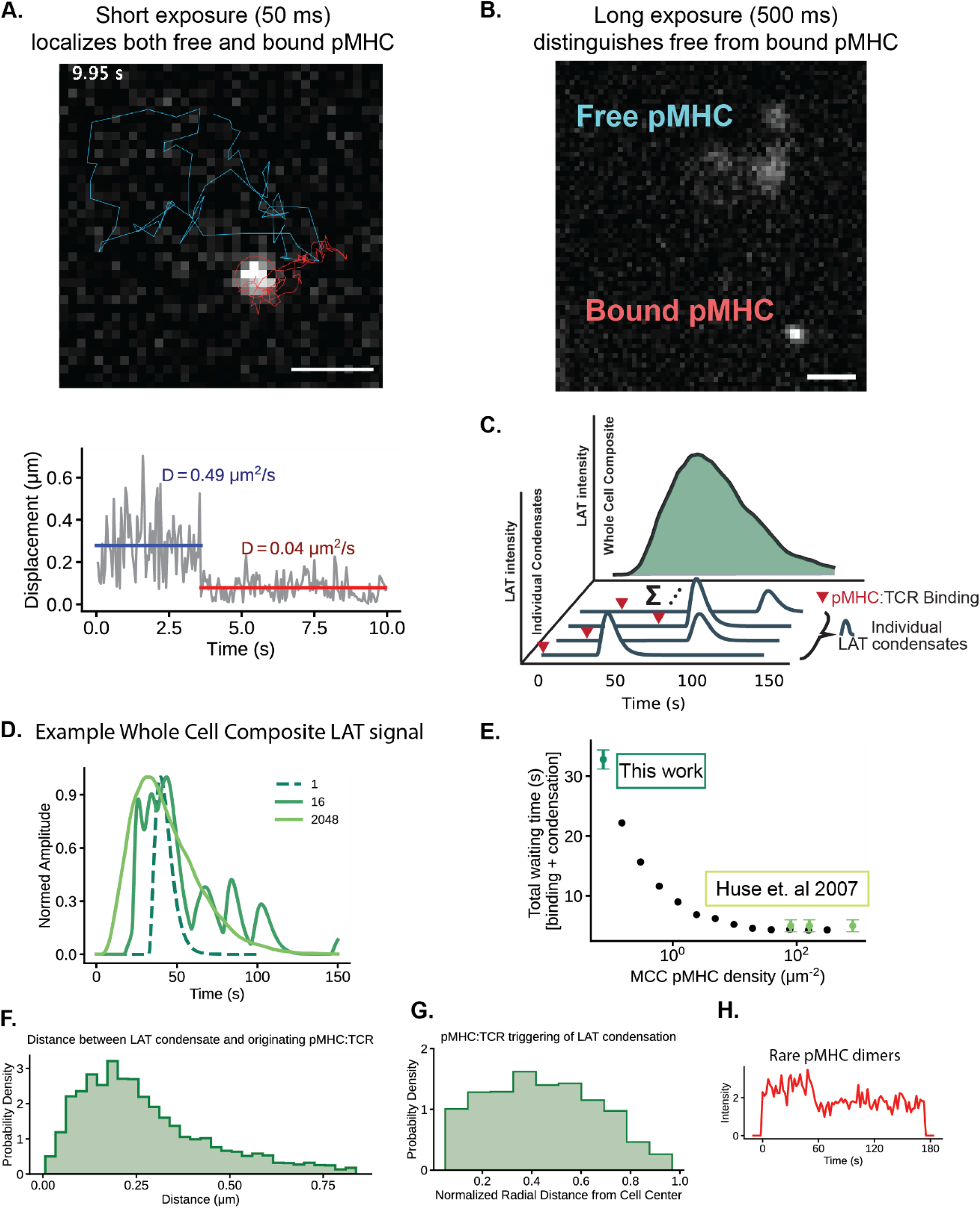
(A) Top: TIRF image showing the full trajectory of a binding event. The MCC(Atto647) pMHC molecule was tracked with 50 ms camera exposure prior to binding (blue) and after binding (red). Bottom: Plot showing the displacement of the above pMHC through time. There is a sudden transition from a free state (blue) to a bound state (red) at 3.5 s, with an accompanying transition in the diffusion coefficient. (B) TIRF image using long camera exposure (500 ms) to distinguish bound pMHC from free pMHC. (C) Schematic showing the integrated LAT response at the cellular level under high stimulation conditions. (D) Example traces showing whole cell LAT response for different numbers of simulated LAT condensate pulses. The dashed line is the average expected single pulse: ≈ 7 s delay from initial cell-bilayer contact until the pMHC binds to the TCR, then a delay time of ≈ 25 s between binding and condensation, and then a LAT condensate lifetime with a lifetime of ≈ 25 s. The solid lines are generated from summing N randomly sampled LAT condensate pulses, renormalized such that the peak amplitude is 1. (E) Plot showing the mean total offset time, from exposure to pMHC ligand until detected LAT signal, for composite cell-wide LAT intensity traces as a function of bilayer density. (F) Histogram of distances between the centroid of the LAT condensate and the centroid of its associated pMHC:TCR binding event. Distances are aggregated from all time frames in which both the condensate and its pMHC:TCR are present, yielding a temporally global distance histogram. (G) The probability of LAT condensation as a function of normalized radial distance 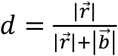, where 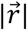 is the distance from the center of the cell to the point of condensation and 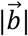 is the distance from the point of condensation to the nearest boundary point of the cell. The distribution was normalized to the area of each annulus corresponding to a particular radial bin. (H) Intensity trace showing the binding of a pMHC dimer. Intensity was normalized to a single step. This class of binding event was excluded from further analysis, i.e., only the effects of monomeric pMHC were studied in detail.

**Figure S2.**
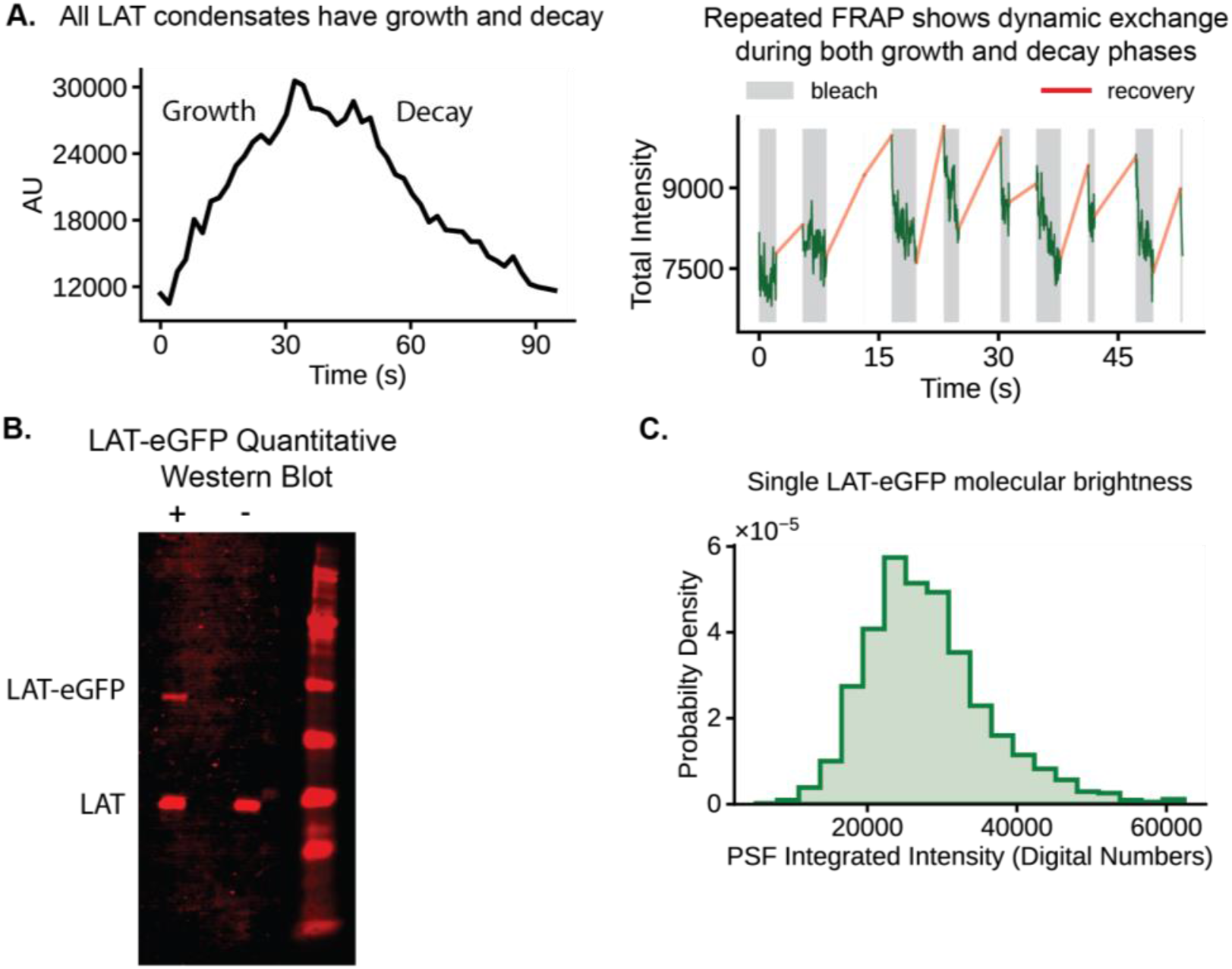
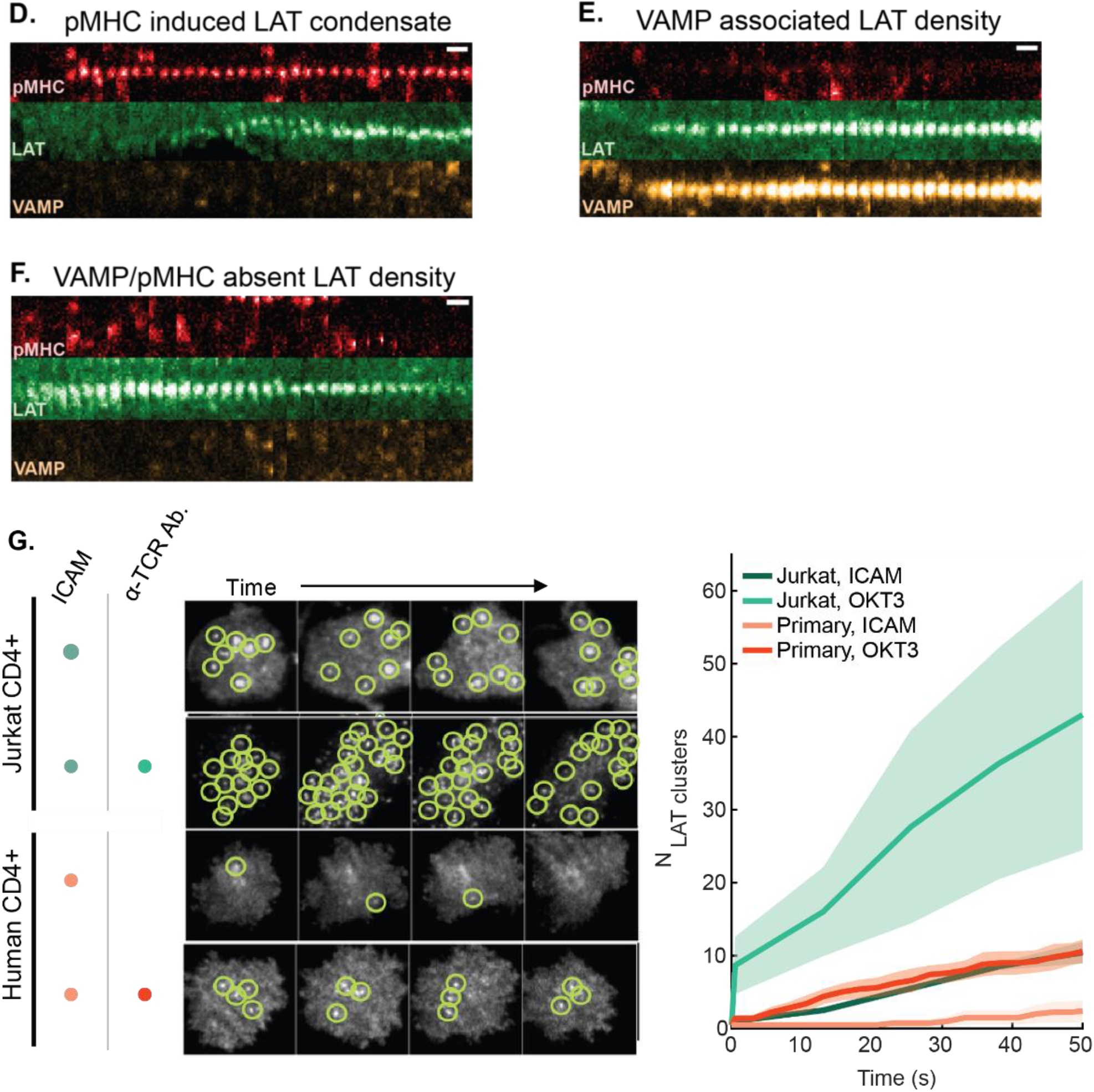
(A) Left: Example time trace of the intensity of LAT condensate, showing the characteristic growth and decay phases intrinsic to all LAT condensates. Right: Periodic bleaching of a LAT condensate shows rapid recovery throughout the lifetime of the LAT condensate. (B) Quantitative western blot using a primary pan-LAT antibody and a secondary IR conjugated antibody. The + lane contained primary T cells transduced with MSCV plasmid containing LAT-eGFP, while the − lane contained untransduced primary T cells. Contrast and background subtraction have been applied to this image. FACS showed %15.2 of the cells in this batch were successfully transduced with LAT-eGFP. (C) Distribution of the integrated net intensity within spot detections of single LAT-eGFP molecules at the plasma membrane, used for quantitative fluorescence to determine the number of LAT molecules within LAT condensates; see **Movie S7**. (D - F) Representative time-series of TIRF images of primary murine T cells expressing LAT-mScarleti and VAMP-mNeonGreen deposited onto supported membranes containing ICAM-1 and MCC(Atto647) pMHC:TCR. (D) Condensation of LAT in response to a pMHC:TCR binding event, but lacking any notable VAMP dynamics. (E) A LAT density associated with VAMP, but not MCC. (F) A LAT density either associated with dark MCC binding, or has an alternate origin, independent of VAMP and MCC. (G) Left: Time series TIRF images of LAT clustering seen in both Jurkat cells (top) and primary human T cells (bottom) under adhesion only (human his-ICAM) and adhesion + stimulation conditions (biotin-OKT3). Bilayers contained biotin-CAP-PE and Ni-NTA-DOGS lipids; and were coated with streptavidin and activated with Ni^2+^ to allow binding with biotin-OKT3 and his-ICAM-1, respectively. Right: plot showing the cumulative numbers of distinct LAT condensates seen through time. Compare to Figure 5A.

**Figure S3.**
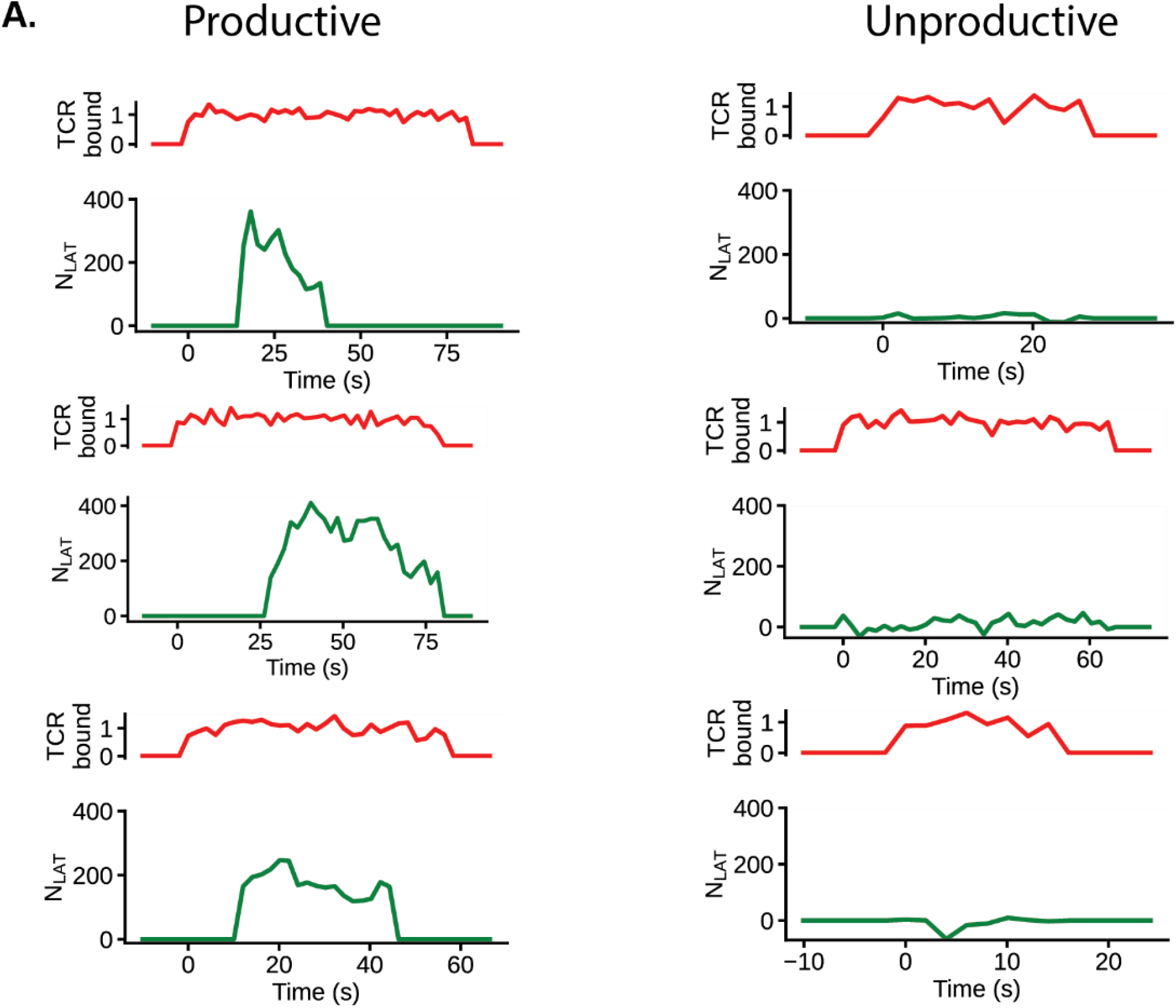
(A) Representative traces of monomeric pMHC:TCR binding and its associated LAT dynamics, compare to Figure 1H. Red traces are the normalized intensity of the bound pMHC. Green traces follow the local maxima of LAT intensity near the pMHC binding event, remapped to number of LAT (see Figure 2).

**Figure S4.**
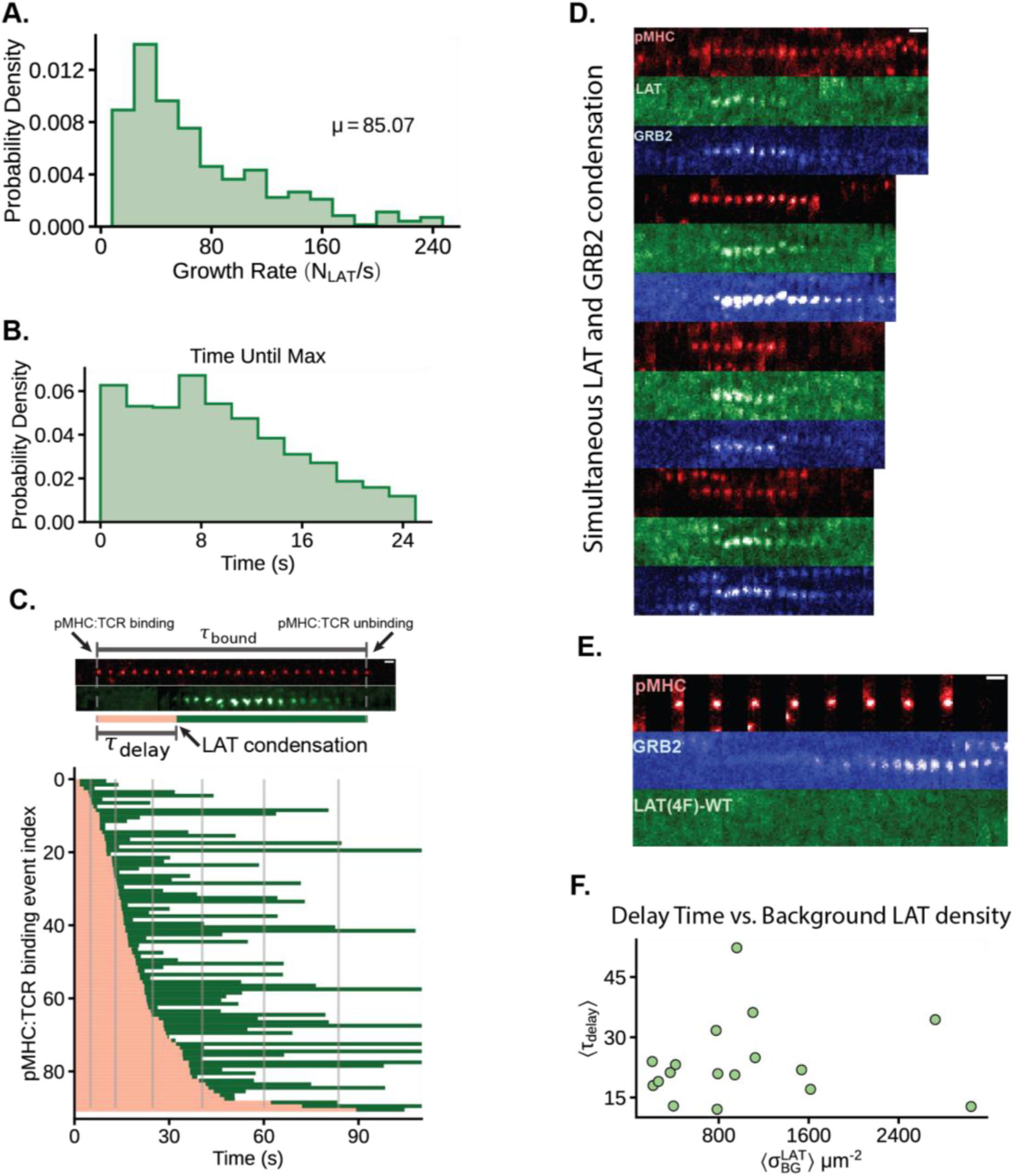
(A) Histogram of the growth rate of LAT condensates within the first 4-6 s of detection. (B) Histogram of the time it takes for LAT condensates to reach peak net intensity. (C) Ensemble of 91 pMHC:TCR binding events that produce LAT condensates illustrating the distribution of LAT delay times. (D) More representative TIRF images of LAT-eGFP(P2A)GRB2-mCherry examples showing the strong coincidence of LAT condensation and GRB2 clustering. (E) Representative trace of a cell expressing LAT(4F)-eGFP(P2A)GRB2-mCherry showing that LAT(4F) fails to participate in pMHC triggered clustering events. (F) Scatter plot of the mean LAT condensation delay time observed in a cell against the mean background LAT density within the same cell. Each point is a cell average.

**Figure S5.**
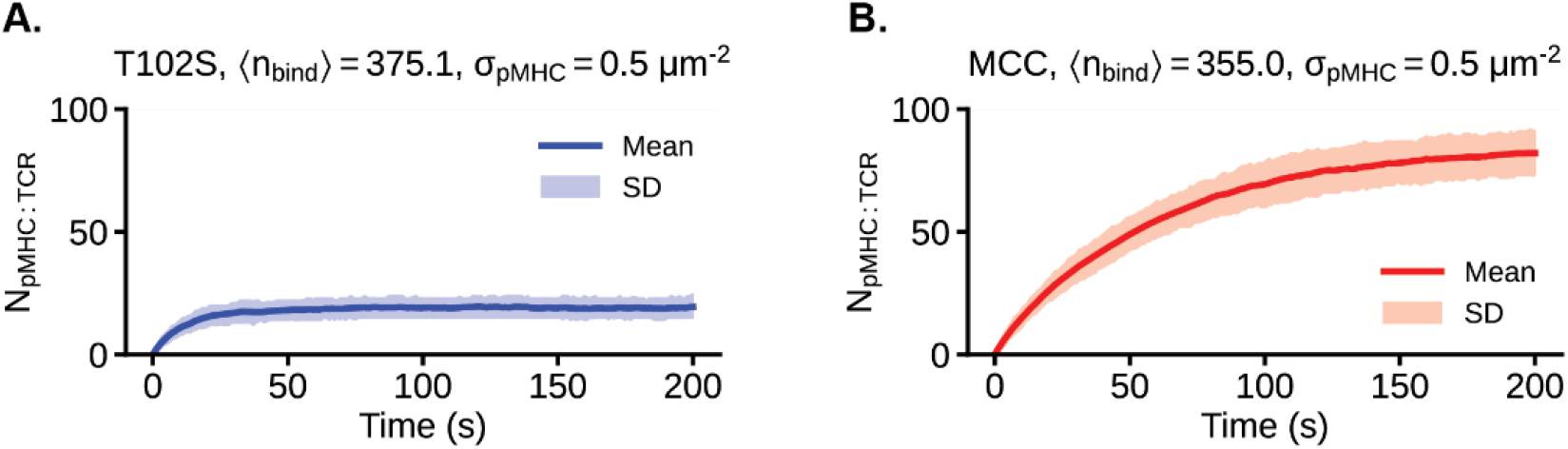
(A) Plot of the number of pMHC:TCR complexes through time for T102S pMHC. From 250 Gillespie simulated trajectories: the mean along with the middle 68% of trajectories (SD) were plotted. (B) Plot of the number of pMHC:TCR complexes through time for MCC pMHC. From 250 Gillespie simulated trajectories: the mean along with the middle 68% of trajectories (SD) were plotted. See Methods for more details.

**Movie S1** Histidine tagged MHC loaded with MCC(C) peptide labelled with Atto647 diffuses on an Ni-NTA lipid bilayer and transitions to a bound state underneath a T cell. ICAM-1 is present for cellular adhesion.

**Movie S2** A primary murine T cell expressing TCR(AND) and LAT-eGFP is deposited onto a bilayer containing 15.9 μm^−2^ of MCC pMHC. ICAM-1 is present for T cell adhesion.

**Movie S3** A primary murine T cell expressing TCR(AND) and LAT-eGFP is deposited onto a bilayer containing 0.43 μm^−2^ of MCC pMHC. ICAM-1 is present for T cell adhesion.

**Movie S4** Zoomed in example of LAT condensation near a pMHC:TCR binding event. A primary murine T cell expressing TCR(AND) and LAT-eGFP is deposited onto a bilayer containing MCC(Atto647) pMHC. Using a camera exposure of 500 ms on the 640 nm channel, bound pMHC were selectively visualized. LAT (green) is shown to condense near a MCC pMHC (red) binding event.

**Movie S5** Cell wide behavior of LAT condensation relative to pMHC:TCR binding events. A primary murine T cell expressing TCR(AND) and LAT-eGFP is deposited onto a bilayer containing MCC(Atto647) pMHC. Using a camera exposure of 500 ms on the 640 nm channel, bound pMHC were selectively visualized. LAT (green) is shown to condense near MCC pMHC (red) binding events.

**Movie S6** An example image of LAT-mCherry condensation. The condensation behavior showed no noticeable qualitative difference from LAT-eGFP. Primary murine CD4+ T cells transduced with LAT-mCherry were deposited onto supported lipid bilayer with ICAM and MCC(Atto647N)-MHC (0.56 μm^−2^). LAT-mCherry condensation was observed over time with TIRF microscopy (560 nm excitation). Note that this measurement was conducted with calcium-responsive dye for another experiment (data not shown), but LAT-mCherry behavior was not affected by the dye.

**Movie S7** Example of a cell with low expression of LAT-eGFP. A primary human T cell expressing LAT-eGFP was imaged continuously with 50 ms camera exposure. Spots that underwent single step photobleaching were used to construct a molecular brightness histogram.

**Movie S8** Multichannel monitoring of MCC pMHC (red, left), LAT-eGFP (green, middle), and NFAT (pink, right) within a primary murine T cell. T cells expressing LAT(G135D)-eGFP were deposited onto bilayers containing MCC(Atto647) pMHC and ICAM-1.

## METHODS

### Lead Contact

Further information and requests for resources and reagents should be directed to and will be fulfilled by the Lead Contact, Jay Groves.

### Materials Availability

Recombinant protein and DNA used in this study may be solicited from the Lead Contact.

### Data and Code Availability

An example raw microscopy acquisition is available in the supplemental data as raw.zip and was used to make to Movie S4 and Movie S5. Associated metadata is also included.

Code was custom built for the data obtained within this study (unless otherwise stated). Data and code may be solicited from the Lead Contact.

### Experimental Model and Subject Details

#### T cell culture and transduction

The mice used in this study were the cross of (B10.Cg-Tg(TcrAND)53Hed/J)x(B10.BR-H2k2 H2-T18a/SgSnJ) from Jackson Laboratory. All animal work was performed with prior approval by the IACUC committee, Lawrence Berkeley National Laboratory Animal Welfare and Research Committee, under the approved protocol 17703. Lymph nodes and spleens of 6-to 20-week-old mice were stimulated with 1 μM MCC peptide in vitro. Both male and female mice were used within this study and were monitored for clean health prior to organ harvest. Activated T cells were retrovirally transduced using Platinum-Eco cell– derived supernatants. Platinum-Eco cells (Cell Biolabs, San Diego, CA) were transfected with the desired plasmid using linear, polycationic polyethylenimine (Sigma-Aldrich). After 48 hours, retrovirus-containing supernatant was used to spinfect T cells in the presence of polybrene (4 mg/ml) on day 3 after activation. Typical transduction efficiencies ranged from 10-30%. The entire population was used for imaging experiments; 0.5 million to 2 million cells were deposited onto each bilayer. Positive fluorescent cells were easily distinguished from negative cells. Overall phenotypic behavior of crawling and spreading were comparable between the two populations using RICM to ensure transduction of the various constructs did not alter basic T cell behavior.

### Method Details

#### Protein Purification

Histidine-tagged MHC class II I-E^K^ and ICAM-1 were expressed and purified as previously described (Nye and Groves, 2008). MHC II with C-terminal hexahistidine tags on both α and β chains were expressed using a baculovirus expression system in S2 cells and purified using a Ni–nitrilotriacetic acid (NTA) agarose column (Qiagen). The histidine-tagged MHC bacmid (Malherbe et al., 2004) was a gift of L. Teyton (Scripps Research Institute) and M. Davis (Stanford University). The bacmid for ICAM-1 with a C-terminal decahistidine was synthesized, and it was similarly expressed and purified in High Five cells (Invitrogen) (Nye and Groves, 2008)

#### DNA constructs

The murine stem cell virus (MSCV) backbone was used (similar to AddGene #91975) for retroviral transduction of primary murine T cells. Plasmids were constructed using Gibson Assembly and cloned using XL1-BLUE (Agilent Technologies) E. coli.

MSCV-NFAT-mCherry was constructed from a plasmid containing a GFP fusion to the regulatory domain of the murine NFAT1 (NFATc2) protein in a murine stem cell virus vector (Marangoni et al., 2013) and was a gift of F. Marangoni (Harvard Medical School). This truncated form of NFAT1 [pMSCV-NFAT1(1–460)-GFP] contains the regulatory domain that controls the nucleocytoplasmic shuttling of NFAT but lacks the DNA binding domain. A second version of the plasmid was generated to replace the GFP coding sequence with mCherry.

MSCV-mNeonGreen-GRB2 was constructed from a plasmid containing human GRB2 that was a gift from Neal Shah (Lo et al., 2019). An mNeonGreen license and plasmid stock was purchased from Allele Biotechnology.

MSCV-LAT-eGFP was constructed from a plasmid containing cDNA for Mus Musculus Linker for Activation of T cells (NM_010689.3).

MSCV-LAT-eGFP-P2A-mNeonGreen-GRB2 was subcloned from the constructs above with the addition of the P2A sequence that was ordered as an oligonucleotide from Elim Biopharmaceuticals, Hayward, CA.

#### Peptide Synthesis and Labelling

Moth cytochrome C 88–103 peptide (MCC88–103; abbreviated as MCC) and previously characterized variants (Corse et al., 2010) were synthesized and lyophilized on campus (D. King, Howard Hughes Medical Institute Mass Spectrometry Laboratory at University of California, Berkeley) or commercially (Elim Biopharmaceuticals, Hayward, CA). A short flexible linker of three amino acids and terminal cysteine was added to the C terminus for fluorophore labeling. The sequences are as follows: MCC (ANERADLIAYLKQATK), MCC(C) (ANERADLIAYLKQATKGGSC), T102S (ANERADLIAYLKQASK), T102S(C) (ANERADLIAYLKQASKGGSC), and T102E (ANERAELIAYLTQAAEK). For dye conjugation, the cysteine-containing peptide sequences were reacted with the maleimide-containing organic fluorophore of interest (Atto647N, Atto565, or Atto488; Atto-Tec GmbH, Siegen, Germany) in phosphate-buffered saline (PBS) with a trace amount of 1-propanol. The labeled peptides were purified using a H_2_O/acetonitrile gradient on a C18 reverse-phase column (Grace Vydac, Deerfield, IL) in the AKTA Explorer 100 FPLC system (Amersham Pharmacia Biotech, Piscataway, NJ). Mass spectrometry was used to confirm the peptide identity after purification.

#### Bilayer Preparation

##### Ni^2+^ chelated bilayers

Glass-supported lipid bilayer membranes were prepared in imaging chambers and functionalized with proteins in a manner similar to prior experiments (Lin et al., 2009; O’Donoghue et al., 2013). At 18 to 24 hours before imaging, MCC and variant peptides were loaded onto MHC II I-E^K^ at 37°C in peptide-loading buffer [PLB – 1% (w/v) bovine serum albumin (BSA) in PBS (pH 4.5) with citric acid]. Just before exposure to bilayers, dye-peptide-MHC complexes in PLB were purified using a 10,000–molecular weight cutoff spin concentrator (Vivaspin 500, GE Healthcare, Pittsburgh, PA) and 1x TBS wash [TBS; 19.98 mM tris and 136 mM NaCl (pH 7.4); Mediatech Inc., Herndon, VA], and resuspended to 300 uL. Small unilamellar vesicles (SUVs) are formed using tip sonication with the composition of 98 mole percent (mol %) 1,2-dioleoyl-snglycero-3-phosphocholine (DOPC) and 2 mol % 1,2 dioleoyl-sn-glycero-3-[(N(5-amino-1-carboxypentyl) iminodiacetic acid) succinyl] (nickel salt) (Ni2+-NTA-DOGS) (Avanti Polar Lipids, Alabaster, AL) in Milli-Q water (EMD Millipore, Billerica, MA), (Lin et al., 2009). In addition, #1.5 25-mm-diameter round coverslips (Thomas Scientific #1217N82) were ultrasonicated in 1:1 isoporopyl/H_2_O and etched for 5 min in piranha solution (3:1 H_2_SO_4_/H_2_O_2_). 35-mm AttoFluor chambers (Fischer Scientific #A7816) were used to house the etched coverslips. A 1:1 mixture of SUVs and 1x TBS was introduced into the chambers, and bilayers were allowed to form through vesicle rupture for at least 30 min. After rinsing, the bilayer was activated with 100 mM NiCl_2_ for 5 min. Samples were exchanged to a live T cell imaging buffer [LCB - 1 mM CaCl_2_, 2 mM MgCl_2_, 20 mM Hepes, 137 mM NaCl, 5 mM KCl, 0.7 mM Na_2_HPO_4_, 6 mM D-glucose, and 0.1% (w/v) BSA]. Immediately before imaging, the bilayer was incubated for 25 min with His-pMHC (≈ 10 pM) and His-ICAM-1 (≈ 10 nM) in LCB. The bilayer was rinsed with imaging buffer after the incubation, and the His-tagged/Ni-NTA– bound proteins were allowed to equilibrate on the bilayer for 20 min. The resulting supported membranes typically display ICAM-1 at 100 to 200 μm^−2^ and pMHC at 0.1 to 2 μm^−2^. The chamber was equilibrated to 37°C, and T cells resuspended in T cell imaging buffer were introduced. All imaging was done in 37°C.

##### OKT3 streptavidin bilayers

SUVs were prepared, ruptured onto coverslips, and activated with NiCl_2_, as above, but with a lipid composition of 98 mol % DOPC, 2 mol % Ni-NTA-DOGS, and 0.02 mol % of 1,2-dioleoyl-sn-glycero-3-phosphoethanolamine-N-(cap biotinyl) (biotin-CAP-PE) (Avanti Polar Lipids, Alabaster, AL). Streptavidin (Sigma-Aldrich, St. Louis, MO) was filtered through a 0.1 μm centrifugal filter (EMD Millipore, Billerica, MA) to remove aggregates, diluted with 1x LCB to 0.5 μg/mL, and then incubated with the bilayers for 45 minutes, rinsed with 1x LBC, and then incubated with biotinylated anti-human CD3 antibody (OKT3) (BioLegend # 317319) at 1 ug/mL for 35 minutes, followed by a rinse with 1x LCB. The bilayers were then incubated with histidine tagged human ICAM-1 (≈ 10 nM) in LCB for 20 minutes, then rinsed with LCB. T cells were introduced as described above.

#### Imaging

All imaging experiments were performed on a motorized inverted microscope (Nikon Eclipse Ti-E; Technical Instruments, Burlingame, CA) with a motorized Epi/TIRF illuminator, a motorized Intensilight mercury lamp (Nikon C-HGFIE), and a motorized stage (MS-2000; Applied Scientific Instrumentation, Eugene, OR). A laser launch with 488-, 560-, and 640-nm diode lasers (Coherent OBIS, Santa Clara, CA) was aligned into a custom-built fiber launch (Solamere Technology Group Inc., Salt Lake City, UT). For TIRF imaging, laser illumination was reflected through the appropriate dichroic beam splitter (ZT488/ 647rpc, Z561rdc with ET575LP) to the objective lens [Nikon (1.47, numerical aperture; 100×), TIRF; Technical Instruments, Burlingame, CA]. RICM and epifluorescent excitation were filtered through a 50/50 beam splitter or band-pass filters (D546/10×, ET470/40×, ET545/30×, and ET620/60×). All emissions were collected through the appropriate emission filters (ET525/50M, ET600/50M, and ET700/75M) and captured on an EM-CCD (iXon 897DU; Andor Inc., South Windsor, CT). All filters were from Chroma Technology Corp. (Bellows Falls, VT). All microscope hardware was controlled using MicroManager (Edelstein et al., 2014).

Laser power was measured at the sample plane with a Newport Power Meter Model 1918-R controller and a 918D-SL-0D2R.

Single-molecule TIRF images are taken at the same power dosage (*power* * *exposure*), either by high laser illumination intensity (8 mW) and short exposure time (25 ms) or by low laser illumination intensity (0.4 mW) and long exposure time (500 ms). Single-molecule TIRF images of long exposure time (500 ms) were collected every 2 to 4 s to localize single pMHC-TCR using a laser illumination intensity of 0.4 mW. LAT-eGFP was monitored at 0.4 mW every 2 to 4 s. When cells were imaged to measure LAT condensation only, the cells were imaged as they landed, or within 60 s of landing, for a duration of 300 s. Cells imaged for NFAT-mCherry was done by epifluorescence using 150-ms exposure time every 50 s at 3 and 6 μmm above the coverslip to monitor NFAT dynamics. NFAT monitoring continued for 30 minutes unless otherwise stated.

Bleach rates for MCC(Atto647) pMHC were determined by incubating pMHC using 1x PBS in an imaging chamber containing a piranha etched glass coverslips (no bilayer) using pMHC volumes that would typically generate 0.1 μm^−2^ pMHC. This creates spatially fixed pMHC. The imaging conditions were set identical to those used for cellular acquisitions. The number of spots within the centrally illuminated region (approximately 350×350 pixels) were tracked through time. Only tracks that appeared in the first frame and were fully immobile were counted. The population of spots through time was fit with an exponential decay. Only the first 150 frames are used for the fit as there tends to be changing decay dynamics at different time scales (likely due to uneven illumination), and typically cells are only imaged for 150-200 frames.

There is a tradeoff between temporal resolution and bleach rate. For the 2 second time-lapse acquisitions that better capture the moment of LAT condensation, a commonly observed bleach rate was 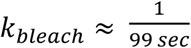, which when compared to the dwell time of MCC, 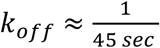, leads to a 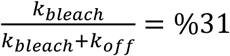 chance that the failure of a pMHC to appear in the next frame was due to bleaching, as opposed to unbinding.

#### Simulation of Whole Cell Integrated LAT Signal

A simple model is constructed to illustrate the effect of sample size on observed delay time, it is not intended to emulate *bona fide* LAT responses. For a static field of potential ligand binders and an already adhered T cell, the “time until a ligand binds” can be modelled to follow an exponentially distribution. In our supported membrane experiments we observe a pseudo-kinetic on-rate of 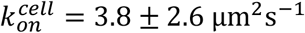 (see *Estimated pMHC on-rate* below). Using this on-rate and a specified ligand density, we can sample when a ligand binds, *t*_*on*_, from an exponential distribution with parameter 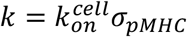. A priori, we expect 0.022 of MCC pMHC binding events with 5cc7 TCR – which has a 6 s dwell time; (O’Donoghue et al., 2013) – to produce a LAT condensate (see *A priori success probability of pMHC binding* below). Of those productive binding events, two random variables were generated, first a random delay time (from the distribution in Figure 4B) and a “random” pulse of LAT signal. A pulse was modelled as a scaled gamma distribution that reflected the shapes of LAT pulses seen in our experiments (compare Figure S1D and Figure S3A).

Sampling from 10s to 1000s of productive binding events, the superposition of the corresponding number of LAT pulses shows that as the number of sample LAT pulses increases, the moment of first detectable signal also becomes earlier (**Figure S1D** and **Figure S1E**).

#### Tracking pMHC and LAT

A combination of the standard TrackMate plugin (Tinevez et al., 2017) for ImageJ and a custom in-house TrackMate plugin was used. The custom plugin was specifically developed to account for the centripetal motion observed by pMHC:TCR binding events. Despite the improved accuracy, each binding event still required manual inspection, using TrackMate’s built-in track inspection tools, to eliminate occasional tracking / detection artifacts. Single-molecule trajectories were verified using single-step unbinding determined by a Bayesian change point detection algorithm (Ensign and Pande, 2010).

A binding event trajectory and dwell time was considered valid if it did not have any branching (i.e. no splits or merges). A LAT delay time was considered valid if its associated pMHC binding event did not have any branching (i.e. no splits or merges).

LAT condensates were initially tracked with TrackMate’s automated workflows and then each track was manually adjusted for proper linking / detection. A pMHC binding event was considered productive if there existed a LAT condensate trajectory whose first frame of detection could be found within 250 nm of the binding event.

### Quantification and Statistical Analysis

Data expressed as *x* ± *y* (SD) represents a mean of *x* and standard deviation of *y*. While (SE) refers to *y* as the standard error of the mean. Data expressed simply as *x* ± *y*, lacking SD or SE, arise from fit parameters, the error of fit was calculated from the square root of the covariance matrix when using the `curve_fit` method in the `scipy` python library. The meaning of the sample size, *n*, is specified in relevant figure legends. Quantification and statistical analysis were done using Python and the following libraries – scikit-image, scipy, pandas, and numpy.

#### Dwell Times

Reported pMHC:TCR dwell times, 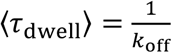, were corrected for photobleaching by measuring the bleach rate of fluorescent pMHC on supported membranes, *k*_bleach_, and then fitting the observed dwell time distribution, *f*_*obs*_(*t*), to the following equation:

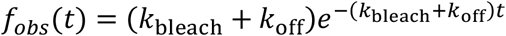

#### FRAP of LAT condensates

Many studies of protein condensation phase transitions use FRAP as a technique to estimate the fluidity of the system (Su et al., 2016; Taylor et al., 2019). A feature of LAT condensates in living cells that complicates this measurement is that the condensates have definite growth and decay phases (**Figure S2A**) as well as centripetal motion towards the center of the cell. However, by intermittently imaging the cell at a fast frame rate (20 ms) and higher laser power followed by multiple seconds to allow recovery, we were able to see significant recovery in the LAT condensates regardless of whether the condensate was actively growing or decaying (**Figure S2B**). Assuming the recovery was entirely reaction-limited, the estimated off-rate for LAT within the condensate was *k*_off_ = 0.24 *s*^−1^, though this is likely a lower-bound estimate.

#### NFAT translocation

NFAT translocation was determined by measuring the fluorescence intensity in the cytosol and nucleus. All images with the masked regions were inspected. The time point at which the mean nuclear NFAT intensity is first detected to be above the mean cytosolic intensity was set as the time point of initial NFAT translocation.

#### Extrapolation from *P*_*LAT*_(*τ*_*pMHC*_)

##### A priori success probability of pMHC binding

The success probability of a ligated TCR (**Figure 3D**), as a function of dwell time, can be used to make a first order approximation of the LAT response to peptides of differing dwell times. This is done by multiplying the dwell time distribution of the pMHC by the success probability function (**Figure 3D**) and integrating over time.

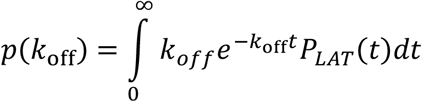

For MCC, which has an exponentially distributed dwell time of ≈ 45 s, the a priori success probability of a binding event producing a LAT condensate is 0.135. Similarly, the a priori success rates for T102S and ER60 binding are 0.044 and 0.003, respectively. Given the same number of spatially isolated binding events, MCC would be expected to produce 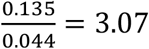 times the number of LAT condensates as T102S. However, experimentally, for an equal density of pMHC, only a 2-fold difference in the number of LAT condensates is observed (**Figure 5A**). Next, we consider potential differences in the number of binding events.

##### Estimated pMHC on-rate

The on-rate of pMHC can change throughout the cell landing and activation process (Pielak et al., 2017). To estimate the on-rate with the presence of photobleaching we consider the simple kinetic scheme:

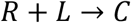

In which receptor (*R*) binds to ligand (*L*) to form a complex (*C*). For a TCR density *σ*_*R*_(*t*), and a *visible* ligand density that exponentially decreases due to bleaching, 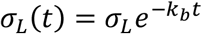, where *σ*_*L*_ is the initial ligand density, then the rate of changing complex density (*σ*_*C*_) is modelled by the following rate equation:

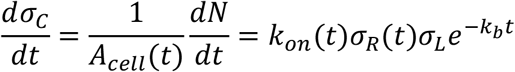

Where *N* is the total number of visible binding events observed over time *t*. To proceed, we instead parameterize a rate equation that uses static mean approximations, and we solve for a pseudo kinetic on-rate, 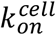 that absorbs the mean cell area and TCR density terms. Integrating over time and solving for 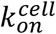 gives:

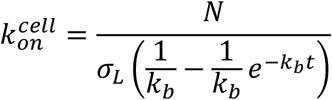

For a collection of 15 T cells expressing TCR(AND), across 3 mice, 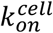 for binding MCC pMHC was determined to be 3.8 ± 2.6 μm^2^s^−1^. T102S had a similar rate.

##### Empirical LAT surplus

T cells expressing TCR(AND) were deposited onto supported lipid bilayers displaying ≈ 600 μm^−2^ ICAM-1 and 0.5 μm^−2^ of pMHC. At this density finding isolated pMHC:TCR is possible, however tracking all pMHC:TCR at this density is difficult due to the numerous merging and splitting events. To test for the presence of depletion effects, and account for photobleaching, a Gillespie simulation (Gillespie, 2007) was performed to estimate the number of binding events over 200 seconds. The parameters used were 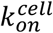 (measured above), the off-rate of the peptide, and the diffusion coefficient of free pMHC (0.55 μm^2^s^−1^). From an ensemble of 250 simulated trajectories, T102S had an average of 375 binding events over 200 seconds (**Figure S5A**), while MCC had an average of 355 binding events (**Figure 5SB**). This suggests that diffusion is fast enough, at this density, to prevent significant depletion effects.

The ratio of empirical to predicted numbers of LAT condensates for MCC was 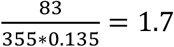; and for T102S the surplus ratio was 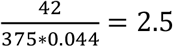.

#### *P*_*LAT*_(*τ*_*MHC*_) and LAT delay times

The combination of dwell time from MCC-MHC:TCR binding and the imaging conditions (power, exposure, etc.) gives rise to an exponential decay of “observed dwell times”.

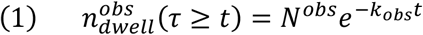

Where *N*^*obs*^ is the total number of observed pMHC:TCR binding events, and *k*_*obs*_ = *k*_*off*_ + *k*_*bleach*_. The empirical frequency distribution for (1) is the orange histogram present in **Figure 3B**. If follows that the observed frequency of productive pMHC:TCR dwell times has the functional form:

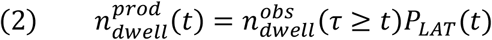

Where *P*_*LAT*_(*t*) is the cumulative probability that an individual pMHC:TCR binding event bound for time *t* has produced a localized LAT condensate. Both (1) and (2) depend on imaging conditions. The empirical frequency distribution for (2) is the red curve present in **Figure 3B**. By dividing 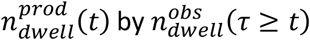, we recover *P*_*LAT*_(*t*) (**Figure 3D**). Moreover, *k*_*obs*_ now merely affects the sampling rate of *P*_*LAT*_(*t*), not the shape of *P*_*LAT*_(*t*).

The histogram of delay times between pMHC:TCR binding and LAT condensation (**Figure 4B**) arises from the following functional form:

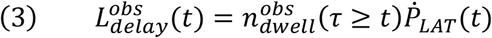

Where 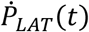 is the probability distribution function of delay time from pMHC:TCR binding until LAT condensates is observed. The number of delay times observed occurring at time *t*, 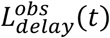, is 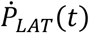 multiplied by the number of binding events that are known to exist for *at least* a duration *t*. As such, (3) is affected by *k*_*obs*_, and for meaningful comparisons of delay time distributions, the imaging conditions and peptide-MHC must be the same, as they are in Figure 4B, 4D, and 4F. What is plotted in **Figure 4B** is 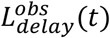 ratioed to the total number of LAT condensates observed to originate from pMHC:TCR binding.

#### *k*_*c*_(*t*) as a Hazard Rate

As derived in the main text, 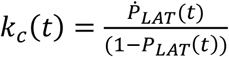. Within the field of survival analysis this quantity is known as a hazard rate, *h*(*t*). Often, one would gather data of “time until failure” for a number of similar objects and create a distribution from this data. The hazard rate, *h*(*t*) is then the probability that a “failure” occurs within time *t* + *dt*, given that it has not yet failed.

For the gamma distribution, the hazard rate *h*^*γ*^(*t*) has a simple analytical expression:

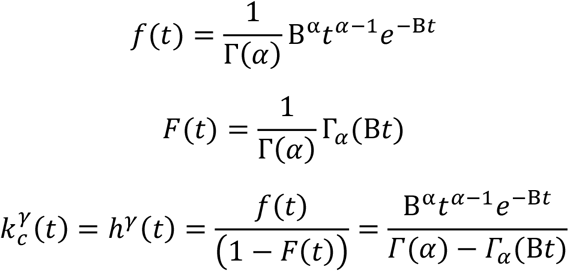

Where Γ(*x*) is the Gamma function, and Γ_*y*_(*x*) is the lower incomplete Gamma function.

The often-simplified form of kinetic proofreading that assumes equally slow steps has a gamma distribution of “success times” (instead of failure times). The first 13 s of the empirical *k*_*c*_(*t*) is used to fit the hazard rate of three different gamma distributions (with one, two, or three equally slow steps). And the result is seen in Figure 3E.

#### Quantitative Fluorescence

To estimate the number of LAT-eGFP molecules within a LAT condensate we first calculate the number of fluorescent LAT molecules, 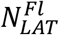, by determining the mean integrated intensity of *single* LAT-eGFP molecules on the plasma membrane (*S*_*I*_). This can be done either by finding a low-expressing cell or bleaching the cell down to single molecule densities and ensuring to analyze those particles with single step photobleaching. Ideally, we could divide the cluster intensity (*C*_*I*_) by the mean PSF intensity:

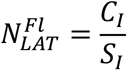

However, there are a few caveats. First, it is often the case that *S*_*I*_ is imaged at different power, exposure, and EMCCD gain conditions than for bulk clusters *C*_*I*_. This is done to stay within the dynamic range of the camera at both expression levels.

Fortunately, for many imaging systems, linear changes in power, exposure, and gain result in linear changes in intensity, but this must be verified. For example, the Andor iXon887 feature RealGain™ technology allows linear inference in the number of incident photoelectrons from fluorescent photons. And LED lasers allow for precise control of power input.

Once linearity in power, gain, and exposure is established then the following normalization will account for the shifting intensity values recorded under different conditions:

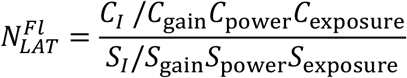

For example, a given cluster intensity *C*_*I*_ recorded at *C*_power_ = 2 will represent half the number of molecules if the same intensity value is recorded with *C*_power_ = 4.

The second caveat is to consider how much of the cellular background LAT is participating within the cluster. If there exists an equal density of free LAT within the cluster region as there does outside the cluster region, then one should use the net cluster intensity to determine the number of extra molecules that exist within the cluster region. However, if the density of free LAT within the cluster region is zero, then the total intensity of the cluster should be used to calculate the number of clustered molecules.

In practice, we used the midpoint between the net intensity and the total intensity.

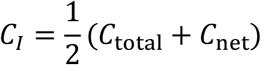

Next, quantification of the western blot will reveal the ratio of exogenous to endogenous LAT (Figure S2F), 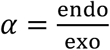. In this study, *α* was determined to be 0.60, and the resulting expression for total LAT is:

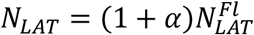

## Notes

### Competing Interest Statement

The authors have declared no competing interest.

