## Supplementary material for "Discrete LAT condensates encode antigen information from single pMHC:TCR binding events": Movies: notes.txt

The power at objective is 0.22 that of the "set-point" power indicated in the acquisition titles. The original acquisition titles can be found in the metadata under the key "prefix".

The bilayer had MCC(Atto647) loaded into his-MHC. Also approximately 300 molecules / um^2 of his-ICAM-1.

This aqcuisition had 1 mW of setpoint power for the 640 channel and 0.5 mW of setpoint power for the 488 channel, and 100 EMCCD gain on the 488 channel.
